# Rational reprogramming of cellular states by combinatorial perturbation

**DOI:** 10.1101/468249

**Authors:** Jialei Duan, Boxun Li, Minoti Bhakta, Shiqi Xie, Pei Zhou, Nikhil V. Munshi, Gary C. Hon

## Abstract

Ectopic expression of transcription factors (TFs) can reprogram cell state. However, due to the large combinatorial space of possible TF cocktails, it remains difficult to identify TFs that reprogram specific cell types. Here, we develop Reprogram-Seq to experimentally screen thousands of TF cocktails for reprogramming performance. Reprogram-Seq leverages organ-specific cell atlas data with single-cell perturbation and computational analysis to predict, evaluate, and optimize TF combinations that reprogram a cell type of interest. Focusing on the cardiac system, we perform Reprogram-Seq on MEFs using an undirected library of 48 cardiac factors and separately on a directed library of 10 epicardial-related TFs. We identify a novel combination of 3 TFs which efficiently reprogram MEFs to epicardial-like cells that are transcriptionally, molecularly, morphologically, and functionally similar to primary epicardial cells. Reprogram-Seq holds promise to accelerate the generation of specific cell types for regenerative medicine.

## Introduction

Ectopic expression of transcription factors (TFs) can reprogram cellular state. For example, forced expression of *MyoD1* alone reprograms fibroblasts to myoblasts (*1*). In addition, transduction of *Oct4*, *Sox2*, *Klf4*, and *Myc* converts mouse embryonic fibroblasts (MEFs) to an induced pluripotent stem (iPS) cell state (*2*). This seminal study required three time- and labor-intensive features: 1) an extensive prior knowledge base to inform selection of the original 24 TF pool, 2) generation of a knockin mouse to report on pluripotency (Fbx15-bGeo), and 3) laborious “N-1” TF evaluation that required multiple iterative rounds of screening. Follow up studies have used similar strategies to optimize iPS reprogramming efficiency or identify TF cocktails to reprogram other cell states. For example, *Gata4*, *Hand2*, *Mef2c*, and *Tbx5* reprograms fibroblasts into cardiomyocyte-like cells (*3*, *4*). However, efficiently generating a multitude of cell types by direct reprogramming will require new approaches for systematic TF identification, reprogramming evaluation, and cell-type assessment. To address this problem, several computational approaches have recently been described (*5*–*8*). However, empiric experimental evaluation is not a component of these methods. As the rules governing cellular reprogramming are poorly defined, integrating both computational and experimental observations to identify TF cocktails capable of reprogramming specific cell types could be a viable alternative strategy.

Here, we describe Reprogram-Seq, an approach that leverages organ-specific cell atlas data with single-cell perturbation and computational analysis to predict, evaluate, and optimize TF combinations that reprogram a cell type of interest. Focusing on the cardiac system, we demonstrate two orthogonal approaches for reprogramming epicardial-like cells. First, we apply Reprogram-Seq to screen random combinations of a large 48-factor (48F) library for reprogramming potential. Second, in a more focused approach, we establish a single-cell atlas of the P0 mouse heart to subsequently identify 10 candidate TFs (10F) for epicardial reprogramming. Reprogram-Seq analysis on these TFs identifies a novel 3 TF combination (3F) that more efficiently generates cells resembling an epicardial cell state. Importantly, we show that 3F-reprogrammed MEFs resemble genuine epicardial cells morphologically and functionally. Unlike recent approaches relying purely on computational prediction, Reprogram-Seq empirically tests and evaluates thousands of TF cocktails by direct experimental measurement. Furthermore, direct reprogramming has emerged as a promising approach for cellular therapy (*9*), but the generation of specific cell types on demand remains challenging. Thus, Reprogram-Seq can be applied to reprogram cell types defined by single-cell genomics with important implications for regenerative medicine.

## Results

### Single-cell combinatorial reprogramming with Reprogram-Seq

For a given *in vivo* cell type defined by single-cell RNA-Seq (scRNA-Seq), we hypothesize that overexpression of a cocktail of cell-type specific TFs can drive cellular reprogramming towards these cells. To enumerate, test, and identify combinations of TFs that can efficiently reprogram fibroblasts to this cell type, we have developed an approach called Reprogram-Seq (**Figure S1**), which measures one key phenotype of reprogramming: the transcriptome. Briefly, we infect mouse embryonic fibroblasts (MEFs) with a retroviral library of candidate TFs. At high infectivity, each cell expresses multiple exogenous TFs that drive transcriptional reprogramming. This results in a library of perturbed cells, where different cells can express different combinations of exogenous TFs. To phenotypically characterize these cells, we perform scRNA-Seq. For each cell sequenced, we measure its full transcriptome, which we use to identify groups of cells where exogenous TFs have driven transcriptional reprogramming. Importantly, detection of exogenous TFs does not rely on distal barcoding, thus avoiding complications with barcode recombination (*10*–*12*). By scaling reprogramming experiments to the single-cell level, Reprogram-Seq allows a highly parallel search for reprogramming factors. Furthermore, profiling thousands of individual cells using scRNA-Seq allows many distinct combinations of TF cocktails to be tested simultaneously in a single well-controlled experiment.

### Unbiased reprogramming with 48 factors

To test the feasibility of Reprogram-Seq on the cardiac system (**Figure 1A**), we first defined the transcriptional state of P0 mouse hearts. Our analysis of 15,684 single-cell transcriptomes identified known cardiac cell types and their molecular markers, including cardiomyocytes (*Tnnt2*+), cardiac fibroblasts (*Col1a1*+), epicardial cells (*Wt1*+), and macrophages (*Lys6c*+) (**Figure 1B**, **S2A-B**, **Table S1**).

**Figure 1:**
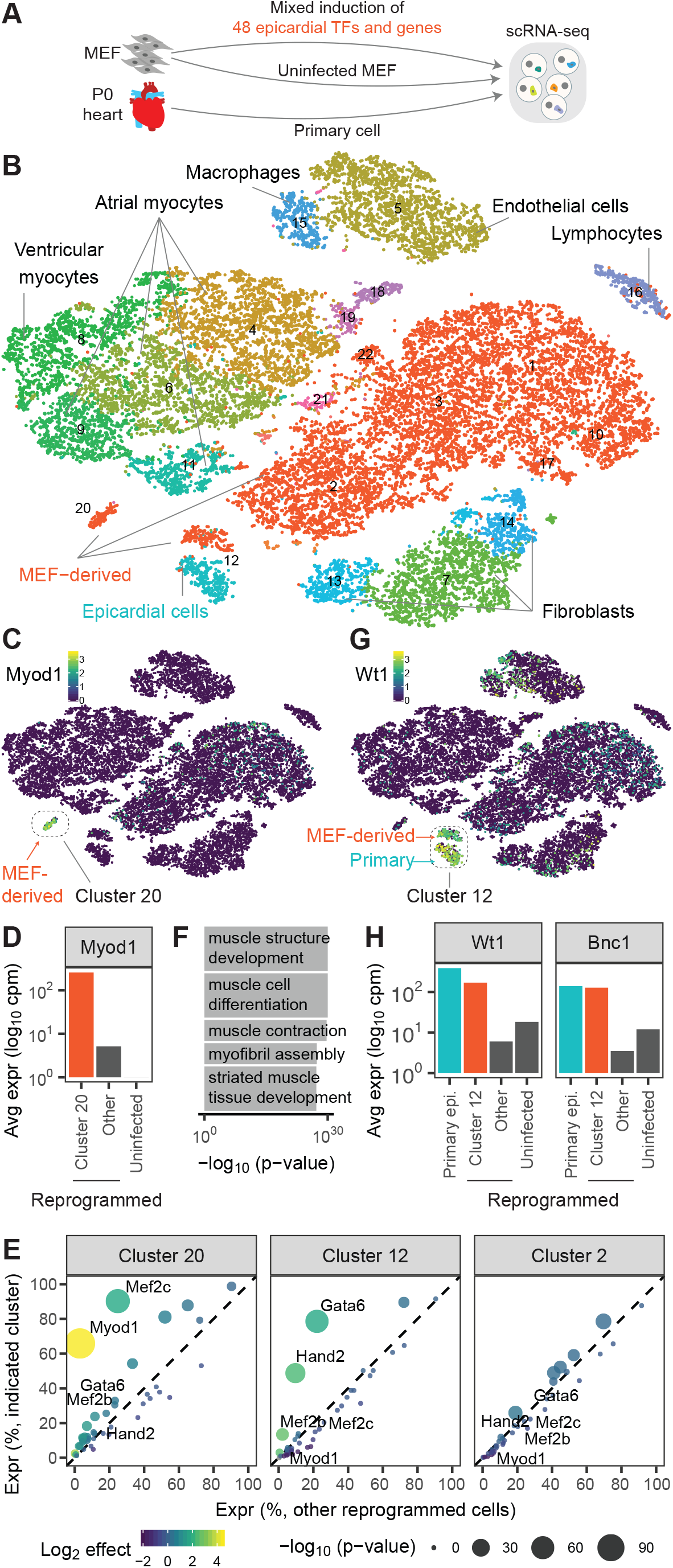
Application of Reprogram-Seq for unbiased reprogramming with 48 factors. A. Experimental design. We performed scRNA-Seq on MEFs reprogrammed with 48F, uninfected MEFs, and primary P0 mouse heart cells. B. 25,776 cells were clustered and plotted using t-SNE (MEF-derived, red; primary cells, non-red). C. Heatmap of cells by *Myod1* expression. MEF-derived cells and Cluster 20 indicated. D. Expression of *Myod1* in MEF-derived cells. E. Differential expression analysis of 48F in MEF-derived clusters, as compared to all other reprogrammed cells. Each dot represents a gene (colored by fold change and sized by p-value). F. Gene ontology analysis for genes differentially expressed in Cluster 20. G. Heatmap of cells by *Wt1* expression. Indicated is Cluster 12, which consists of MEF-derived cells and primary cells. H. Expression of *Wt1* and *Bnc1* in MEF-derived cells.

Next, we tested whether Reprogram-Seq can perform combinatorial TF reprogramming of many TF cocktails at the same time. We created a complex library of 48 TFs and genes (48F) expressed in cardiac cells based on literature curation and bulk transcriptome analysis. Some of these genes have known roles in cardiac biology, including members of the GATA, Hand, Mef2, and T-box families. As a positive control, we also included *MyoD1*, which alone can reprogram fibroblasts to skeletal muscle cells (**Table S2**). We packaged a pooled retroviral library, infected MEFs, and profiled their transcriptomes by scRNA-Seq. The vast majority of infected MEFs clustered together with uninfected MEFs (**Figure 1B**), suggesting that most TFs do not dramatically alter the MEF transcriptome when compared to *in vivo* cell types. However, there were two notable exceptions.

Almost all of the cells in Cluster 20 are MEF-derived and transcriptionally distinct from other MEF-derived cells (**Figure 1B**). This cluster has the highest levels of *MyoD1* expression compared to other MEF-derived cell clusters (**Figure 1C-E**), suggesting that these cells are derived from MEFs infected with retrovirus driving *MyoD1* expression. Supporting this possibility, the genes induced in Cluster 20 are significantly enriched for gene ontology annotations related to muscle development (**Figure 1F**), and the cells in Cluster 20 exhibit endogenous *MyoD1* gene expression (**Figure S3A**). In addition, MEFs separately infected with *MyoD1* retrovirus correlate more with Cluster 20 than with any other cluster (**Figure S3B**). Taken together, these results indicate that *MyoD1* successfully outcompetes 47 other exogenous factors in a subpopulation of cells to push cell state towards a skeletal muscle cell fate, and highlights the fidelity of combinatorial reprogramming achieved by Reprogram-Seq.

Next, we wondered whether Reprogram-Seq could identify MEFs reprogrammed to a cardiac-like cell fate. Interestingly, Cluster 12 contains a mixture of *in vivo* (63.1%) and MEF-derived cells (36.9%) (**Figure 1G**). All of the *in vivo* epicardial cells are found in this cluster, suggesting that the MEF-derived cells may have activated epicardial genes. Consistent with this possibility, the MEF-derived cells in Cluster 12 express known markers of epicardial cells, including *Wt1* and *Bnc1*, which are not in 48F (**Figure 1H**). To identify the exogenous TFs induced in these cells, we examined the expression of 48F. Strikingly, 78.6% of MEF-derived cells in Cluster 12 express *Gata6*, compared with only 22.1% in other MEF-derived cells (**Figure 1E**, **S3C**) (p = 4.4E-62; binomial). Similarly, *Hand2* expression is highly enriched (48.8% Cluster 12; 9.6% other; p = 5.2E-43). These results suggest that *Hand2* and *Gata6* may transcriptionally reprogram MEFs towards an epicardial-like state. Consistent with these findings, mouse genetic studies have previously shown that *Gata6* and *Hand2* function in epicardial development and maintenance (*13*, *14*).

Together, these results indicate that Reprogram-Seq can efficiently search a large combinatorial space to simultaneously identify multiple TF cocktails (*MyoD1* and *Gata6*/*Hand2*) that reprogram MEFs to distinct cell states.

### Rational reprogramming of epicardial cells

Our single-cell analysis of *in vivo* heart cells identified several cell types. While many TF cocktails can drive MEFs towards a cardiomyocyte cell fate, TF cocktails for other cardiac cell types remain limited. Next, we apply Reprogram-Seq to rationally engineer epicardial cells.

To identify a set of candidate TFs for epicardial reprogramming, we performed differential gene expression analysis of *in vivo* epicardial cells (the destination state of reprogramming) compared to MEFs (the origin state). This analysis identified 10 transcription factors (10F) (**Figure 2A**). We were encouraged by the presence of known markers of epicardial cells including *Tbx18* (*15*–*17*), Tcf21 (*18*, *19*), and *Wt1* (*20*, *21*), as well as TFs identified in our previous unbiased reprogramming (*Gata6* and *Hand2*).

**Figure 2:**
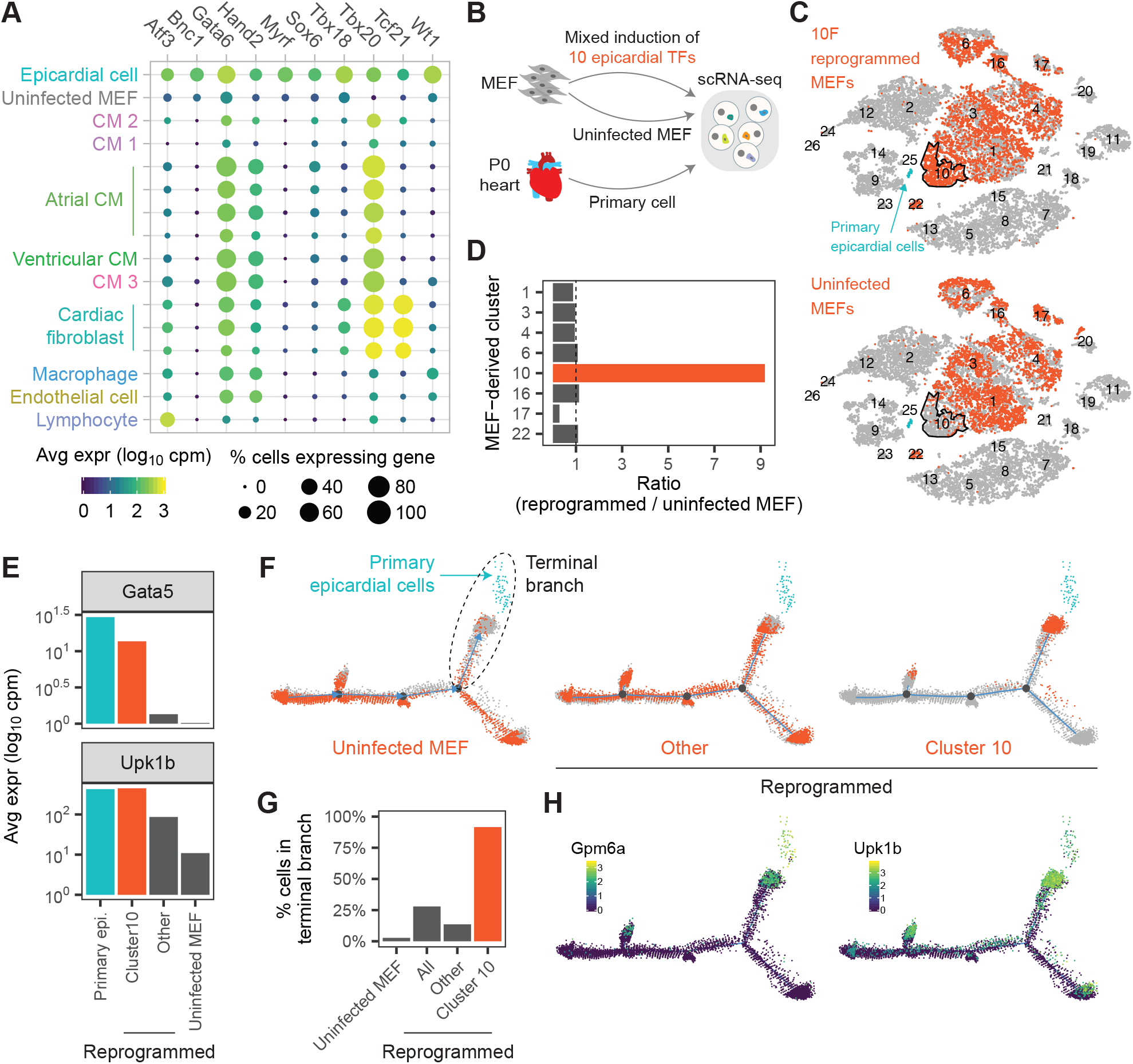
Application of Reprogram-Seq for epicardial reprogramming. A. We identified 10 TFs differentially expressed in primary epicardial cells compared to uninfected MEFs. Shown is the expression of 10F in MEFs and P0 mouse heart cells. B. Experimental design. We performed scRNA-Seq on MEFs reprogrammed with 10F, uninfected MEFs, and primary P0 mouse hearts. C. t-SNE plots consisting of all sequenced cells in (B). (top) 10F-reprogrammed MEFs and (bottom) uninfected MEFs are colored orange. Primary epicardial cells are indicated in blue. Cluster 10 is outlined in black. D. Enrichment of reprogrammed MEFs compared to uninfected MEFs in different cell clusters. E. Expression of epicardial marker genes *Gata5* and *Upk1b* in primary and MEF-derived cells. F. We performed pseudotime analysis on all MEF-derived cells and primary epicardial cells (blue). (left) Uninfected MEFs, (right) Cluster 10 reprogrammed cells, and (middle) other reprogrammed cells are indicated in orange. G. Bar chart indicating the percentage of cells in each group belonging to the terminal pseudotime branch (F). H. Heatmap of cells by (left) *Gpm6a* and (right) *Upk1b* expression.

To test the ability of 10F to convert MEFs to an epicardial-like fate, we performed Reprogram-Seq with 10F after 7 days of retroviral induction (**Figure 2B**). As a control, we also sequenced uninfected MEFs. The vast majority of cells in Cluster 10 (90.0%) are derived from 10F-reprogrammed MEFs (**Figure 2C-D**). To assess if Cluster 10 cells resemble *in vivo* epicardial cells, we examined the expression of known epicardial marker genes *Anxa8*, *Gata5*, *Gpm6a*, *Krt8*, *Krt19*, and *Upk1b* (*15*), none of which are in 10F. We observed significant induction of these epicardial markers in Cluster 10 cells compared with uninfected MEFs and other MEF-derived clusters (**Figure 2E**, **S4A-B**).

To more precisely quantify the efficiency of epicardial reprogramming across the transcriptome, we performed pseudotime analysis across reprogrammed MEFs, control MEFs (origin cell), and primary epicardial cells (target cell). Pseudotime analysis reveals that the starting population of uninfected MEFs is largely distinct from the destination population of primary epicardial cells (**Figure 2F**, left). A critical juncture in this pseudotime is the branch between uninfected MEFs and primary cells, which we denote as the terminal branch. Surprisingly, we observe that the vast majority of reprogrammed cells in Cluster 10 belong to the terminal branch (91.6%), and are particularly enriched in the subset of cells in pseudotime space closest to primary epicardial cells (**Figure 2F-G**). In comparison, only 13.57% of other 10F-reprogrammed MEFs occupy the terminal branch. Consistent with these data, known epicardial markers are highly expressed in terminal branch cells (**Figure 2H**).

### Optimized epicardial reprogramming with 3F

The above results suggest that 10F-reprogrammed MEFs have undergone transcriptional reprogramming towards an epicardial-like state. However, it is possible that only a subset of genes in 10F drives this outcome. To identify subsets of key factors, previous studies have relied on labor-intensive “N-1” approaches, whereby distinct TF cocktails are iteratively tested for reprogramming performance. To accelerate this process, we devised an alternative strategy relying on the single-cell measurements of Reprogram-Seq (see supplemental text). We observed dramatic enrichment of *Atf3*, *Gata6*, and *Hand2* (3F) expression in the reprogrammed cells of Cluster 10, with over half of the cells expressing these factors (**Figure S5A-B**). Collectively, 3F is expressed >2.6-fold more in Cluster 10 than other reprogrammed cells. In addition, we confirmed that cells in Cluster 10 exhibit significant exogenous expression of 3F (**Figure 3A**). These results suggest that 3F may be an optimized TF cocktail for epicardial reprogramming.

**Figure 3:**
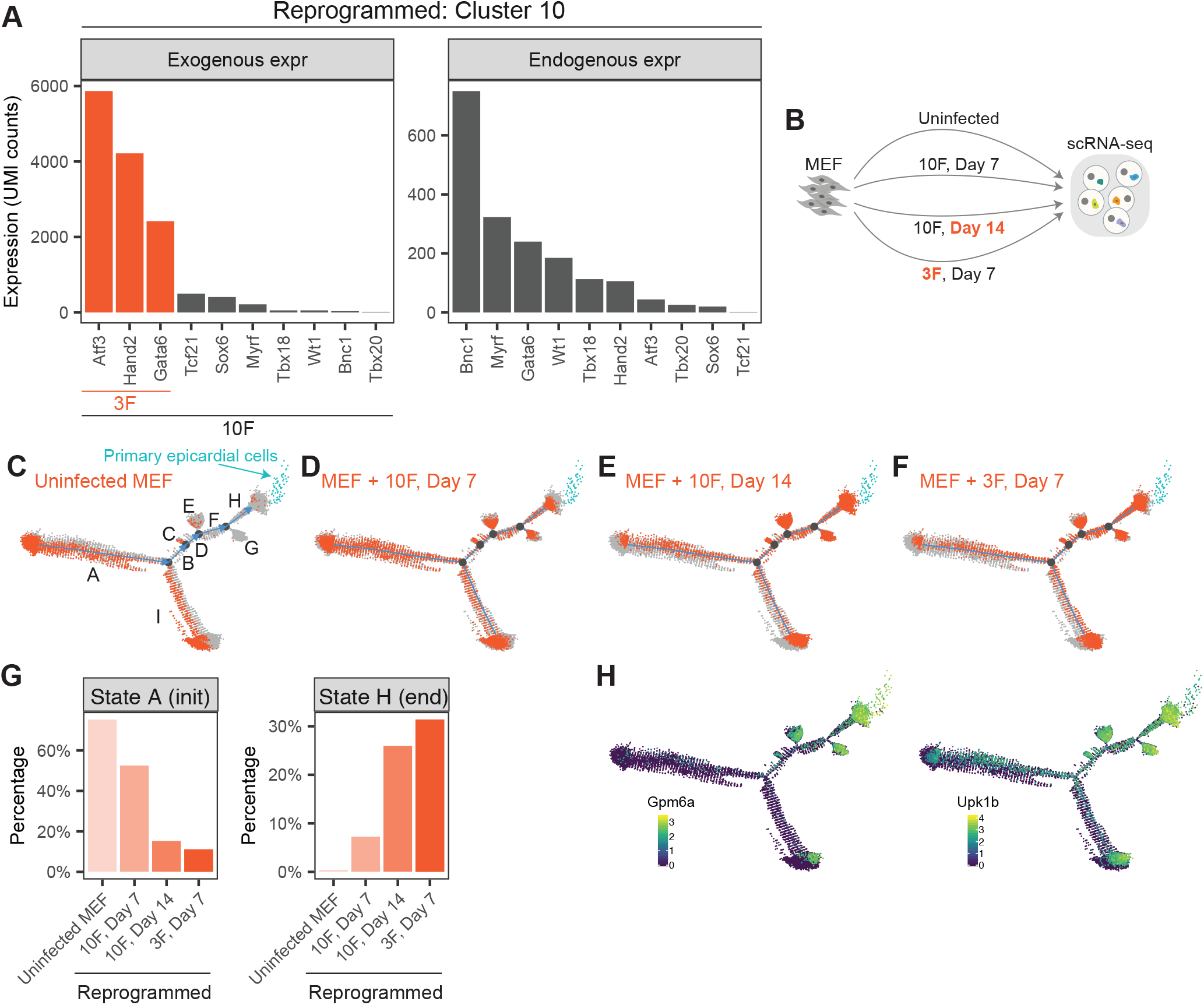
Optimized epicardial reprogramming with *Atf3*, *Gata6*, and *Hand2*. (A) Estimates of exogenous and endogenous expression for 10F in Cluster 10 reprogrammed cells. (B) Experimental design. We performed scRNA-Seq on uninfected MEFs and MEFs reprogrammed with 10F for 7 days, 10F for 14 days, and 3F for 7 days. States A-H of pseudotime are indicated. (C-F) We performed pseudotime analysis on the MEF-derived cells above and primary epicardial cells (blue). Indicated in orange are (C) uninfected MEFs and MEFs reprogrammed with (D) 10F for 7 days, (E) 10F for 14 days, and (F) 3F for 7 days. (G) Bar plot indicating the fraction of cells in each experiment that belong to the (left) initial State A or (right) terminal State H of pseudotime. (H) Heatmap of cells by the expression of the epicardial marker gene (left) *Gpm6a* and (right) *Upk1b*.

To test whether exogenous expression of 3F is sufficient to transcriptionally reprogram MEFs to epicardial-like cells, we performed Reprogram-Seq with 3F for 7 days. To assess performance, we compared 10F reprogramming for 7 days and 14 days (**Figure 3B**), as increased duration of reprogramming has been observed to improve efficacy (*4*, *22*). Combined pseudotime analysis of all cells indicated a trajectory mirroring the previous analysis. Uninfected MEFs were predominantly enriched at the initial pseudotime state A, while all of the primary epicardial cells clustered in the terminal pseudotime state H (**Figure 3C**). Reprogrammed cells occupied states between these two extremes (**Figure 3D-F**). Among 10F cells, we observed a dramatic 3.6-fold (p < 2.2e-16) increase in cells attaining the terminal State H after 14 days (26.0%) of reprogramming compared to 7 days (7.3%), which was accompanied by corresponding decrease in the initial state A (**Figure 3G**, **S5D**). These results confirm that reprogramming efficiency increases with time. Next, we assessed the efficiency of 3F reprogramming. Surprisingly, we found that the performance of 3F for 7 days (31.4% of cells reaching the epicardial state H) exceeded that 10F reprogramming for both 7 days and 14 days. 3F reprogramming was efficient: only 11.2% of cells remained in the initial state A. In addition, 42.3% of cells were on the pseudotime trajectory towards epicardial cells (states BDFH), as indicated by increased expression of the epicardial markers in this path (**Figure 3H**). These observations suggest that 3F is an optimized cocktail for transcriptional reprogramming of MEFs towards epicardial-like cells.

### 3F activates the endogenous epicardial gene regulatory network

To understand the molecular basis of epicardial-like reprogramming, we examined 3F reprogramming along the main pseudotime trajectory consisting of the MEF state A, intermediate states BDF, and epicardial state H (**Figure 4A**). As pseudotime increases along the main trajectory, the expression of 3F increases (**Figure 4B**, **S6A**). This indicates that attaining high expression of exogenous 3F is a key step in epicardial reprogramming, which agrees with previous studies on cellular reprogramming (*23*–*25*). In contrast to the gradual increase of 3F, it is not until the terminal state H that the epicardial marker genes *Myrf*, *Wt1*, and *Bnc1* become endogenously activated (**Figure 4A**).

**Figure 4:**
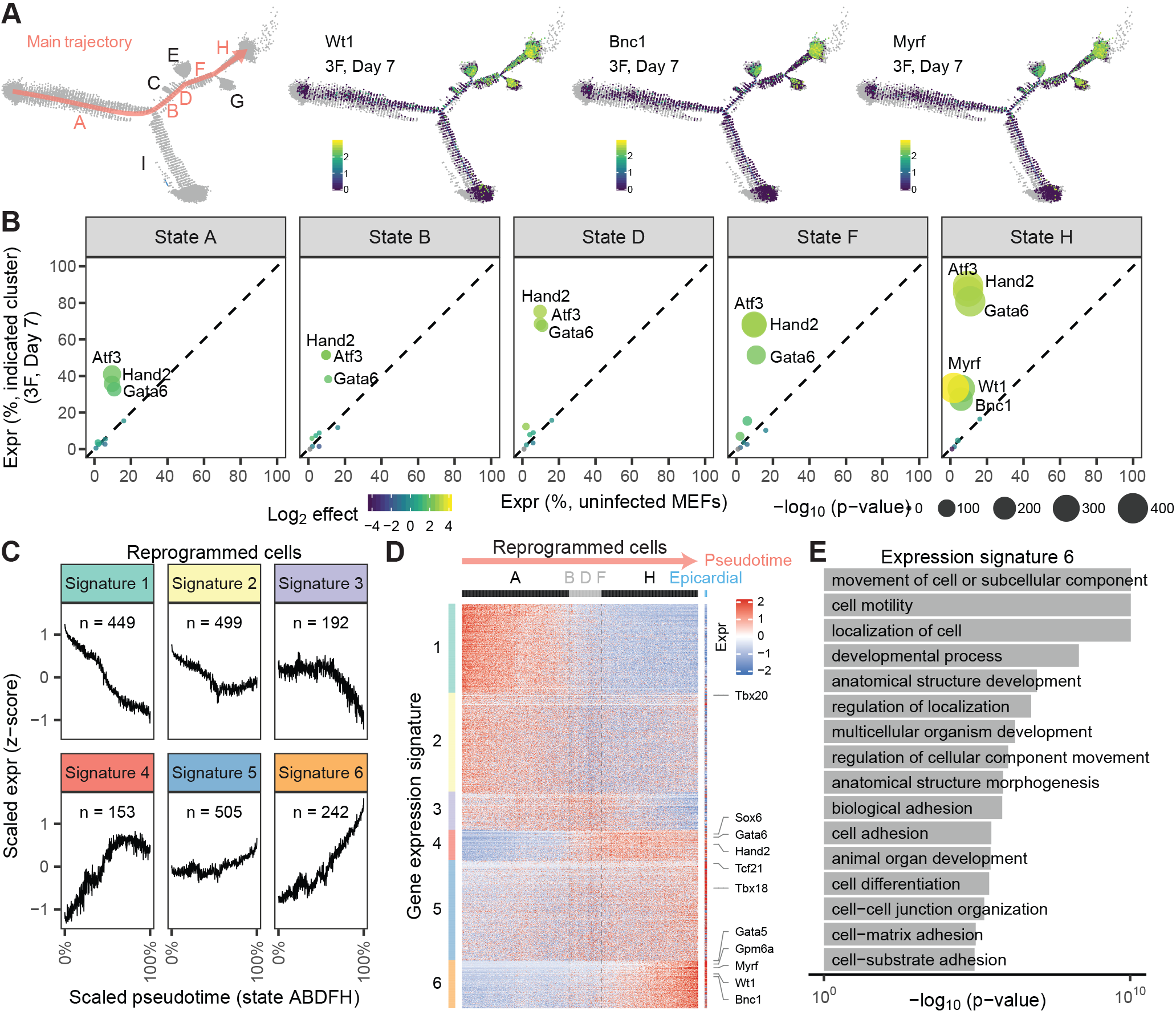
*Atf3*, *Gata6*, and *Hand2* activate the epicardial transcriptional network. A. (left) The main trajectory of pseudotime is highlighted in orange. (middle-right) For MEFs reprogrammed with 3F for 7 days, we plot the expression of epicardial marker genes Wt1, Bnc1, and Myrf. B. Expression of exogenous genes (*Atf3*, *Hand2*, *Gata6*) and endogenous genes (*Myrf*, *Wt1*, *Bnc1*) across pseudotime states in MEFs reprogrammed with 3F for 7 days. C. We identified 2040 genes differentially expressed between MEFs and primary epicardial cells. The genes cluster into six gene expression signatures during pseudo-time. Expression is log10 transformed and Z-normalized. D. Heatmap of the six gene expression signatures. Columns indicate cells ordered by pseudotime (with primary epicardial cells at the far right), rows indicate genes. For visualization purposes, groups of 20 cells are binned. E. Gene ontology analysis of genes with expression signature 6.

To further assess how the MEF gene regulatory network becomes rewired during 3F reprogramming, we clustered genes during pseudotime to identify 6 distinct expression signatures (**Figure 4C**). We observe that 3F significantly reconfigures the expression of ∼2000 genes towards the epicardial state: MEF-specific genes are repressed (Signatures 1-3) and epicardial-specific genes become activated (Signatures 4-6) (**Figure 4D**). Consistent with their intermediate position in pseudotime, the BDF state represents a transitional state of gene expression between states A and H. A few epicardial markers such as *Sox6* are induced early upon commitment to the intermediate BDF branch. Most epicardial markers exhibit the strongest induction late in pseudotime in state H. Indeed, key epicardial genes in Signature 6 (*Gata5*, *Gpm6a*, *Myrf*, *Wt1*, and *Bnc1*) exhibit sustained induction throughout state H, reaching their maximal expression only at the end of pseudotime. Consistent with their roles in epicardial cells, Signature 6 genes are enriched for gene ontology terms relevant to epicardial function including cell motility (p = 9.5e-11) and cell adhesion (p = 3.4e-06) (**Figure 4E**) (*26*). Thus, activation of the endogenous epicardial gene regulatory network occurs late during reprogramming.

Pseudotime also identifies two states (E and G) that branch off the main trajectory. Our analysis indicates that these cells represent alternative cell fates of 3F reprogramming with altered expression of cell cycle genes and signal transduction pathways (see supplemental text) (**Figure S6**).

### Functional validation of epicardial reprogramming

To confirm that Reprogram-Seq has identifies TF cocktails driving conversion of MEFs to epicardial-like cells, we performed reprogramming experiments using traditional bulk approaches and examined several features of epicardial cells (**Figure 5A**). First, we measured the expression of key marker genes by qPCR. For both 10F and 3F, we observe induction of epicardial marker genes *Upk1b*, *Gpm6a*, and *Krt19* (**Figure 5B**, **Table S3**), which are not part of either reprogramming cocktail. Induction of epicardial markers was sustained after 7 and 14 days of reprogramming. Consistent with cell state transition, we also observe that reprogrammed MEFs exhibit decreased expression of the fibroblast marker gene *Col3a1*. Second, microscopy confirms that 10F- and 3F- reprogrammed MEFs adopt a cobblestone morphology which is consistent with that of epicardial cells and which is distinct from the morphology of uninfected MEFs (**Figure 5C**) (*15*). Third, we performed immunostaining for the tight junction marker ZO-1, a molecular marker of epithelial cells such as epicardial cells (*15*). Consistent with epicardial reprogramming, both 10F and 3F cells exhibit ZO-1 protein localize to the cellular periphery (**Figure 5D**). Importantly, reprogramming towards an epicardial cell-like state is specific to 10F and 3F, since GHMT could only generate a-actinin^+^/ZO-1^-^, reprogrammed cells. Fourth, as primary epicardial cells exhibit aldehyde dehydrogenase activity (*15*), we performed the Aldefluor assay to directly measure this activity in reprogrammed cells. Confirming an epicardial-like cell state, we observed nearly 10-fold induction of aldehyde dehydrogenase activity in 10F and 3F cells (**Figure 5E**). Together, these experiments indicate that 10F and 3F can drive the conversion of MEFs towards an epicardial-like cell fate.

**Figure 5:**
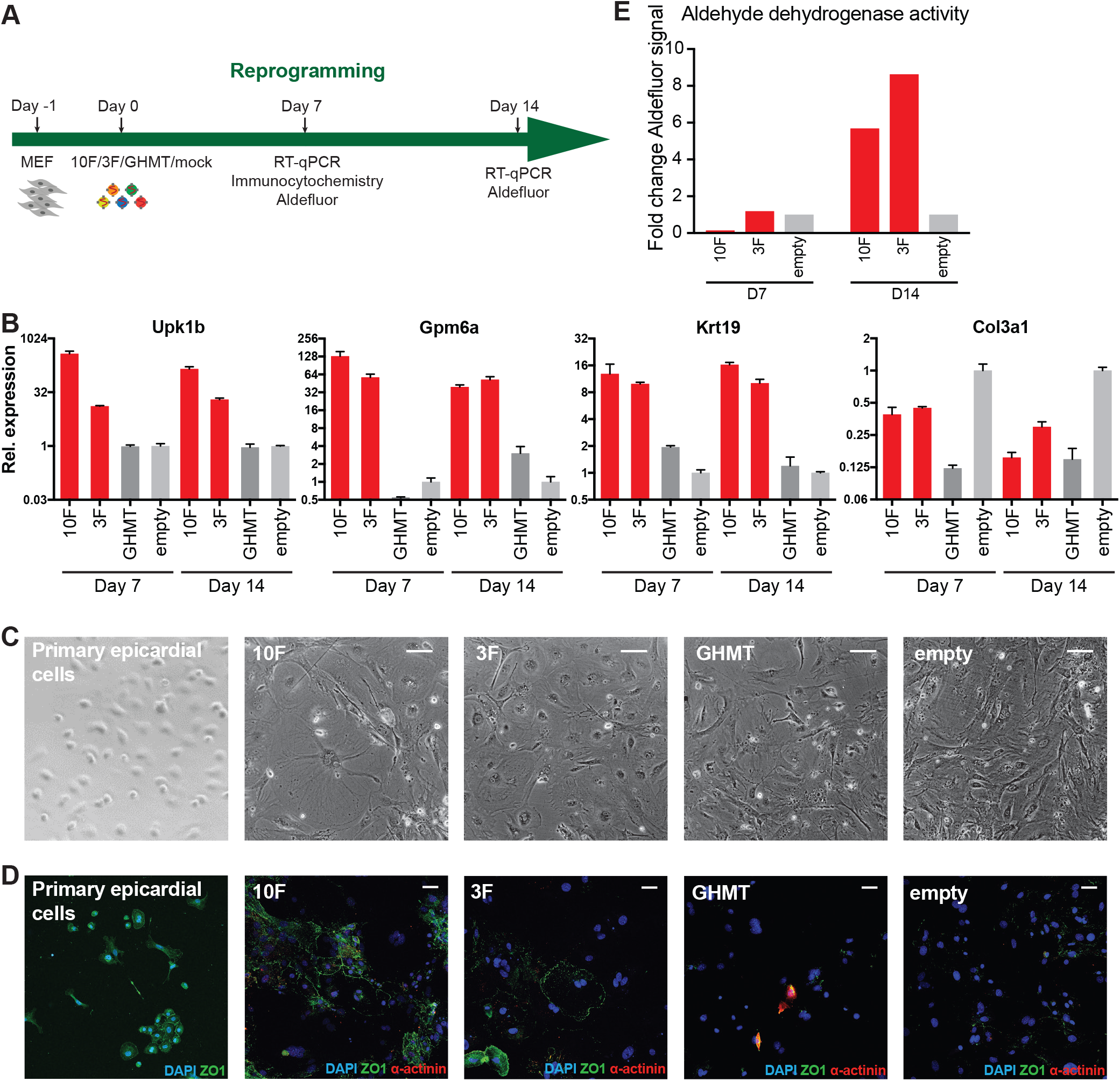
Functional validation of epicardial-like reprogramming. A. Experimental design to functionally validate epicardial-like reprogramming. B. qRT-PCR analysis of epicardial markers (*Upk1b*, *Gpm6a*, *Krt19*) and fibroblast markers (*Col3a1*) in reprogrammed MEFs (10F, 3F, GHMT) and negative control infected MEFs (empty vector). C. Bright field images of primary epicardial cells, uninfected MEFs, and reprogrammed MEFs. Scale bar indicates 100 microns. D. Immunostaining of DAPI, ZO1, and α-actinin in primary epicardial cells and reprogrammed MEFs. Scale bar indicates 50 microns. E. Aldehyde dehydrogenase activity of reprogrammed MEFs as measured by the Aldefluor assay.

### A roadmap for rational cellular reprogramming

We anticipate that Reprogram-Seq can be applied to identify new reprogramming cocktails for cell types defined by cell atlas efforts. The Tabula Muris Consortium has generated scRNA-Seq atlases for 20 organs and tissues spanning ∼60 mouse cell types (*27*). We computationally identify candidate TFs to drive cellular conversion to these cells from MEFs (**Supplemental Table S4**). We find known reprogramming factors for hepatocytes (*Hnf4a*, *Foxa3*) (*28*–*30*) and renal tubular epithelial cells (*Hnf1b*, *Hnf4a*, *Pax8*) (*31*) (**Figure 6A, S7A**). We also identify potential reprogramming factors for cells such as skeletal muscle satellite cells that include TFs with known roles in these cells (*MyoD1*, *Myf5*, and *Pax7*) (*1*) (**Figure 6A**).

**Figure 6:**
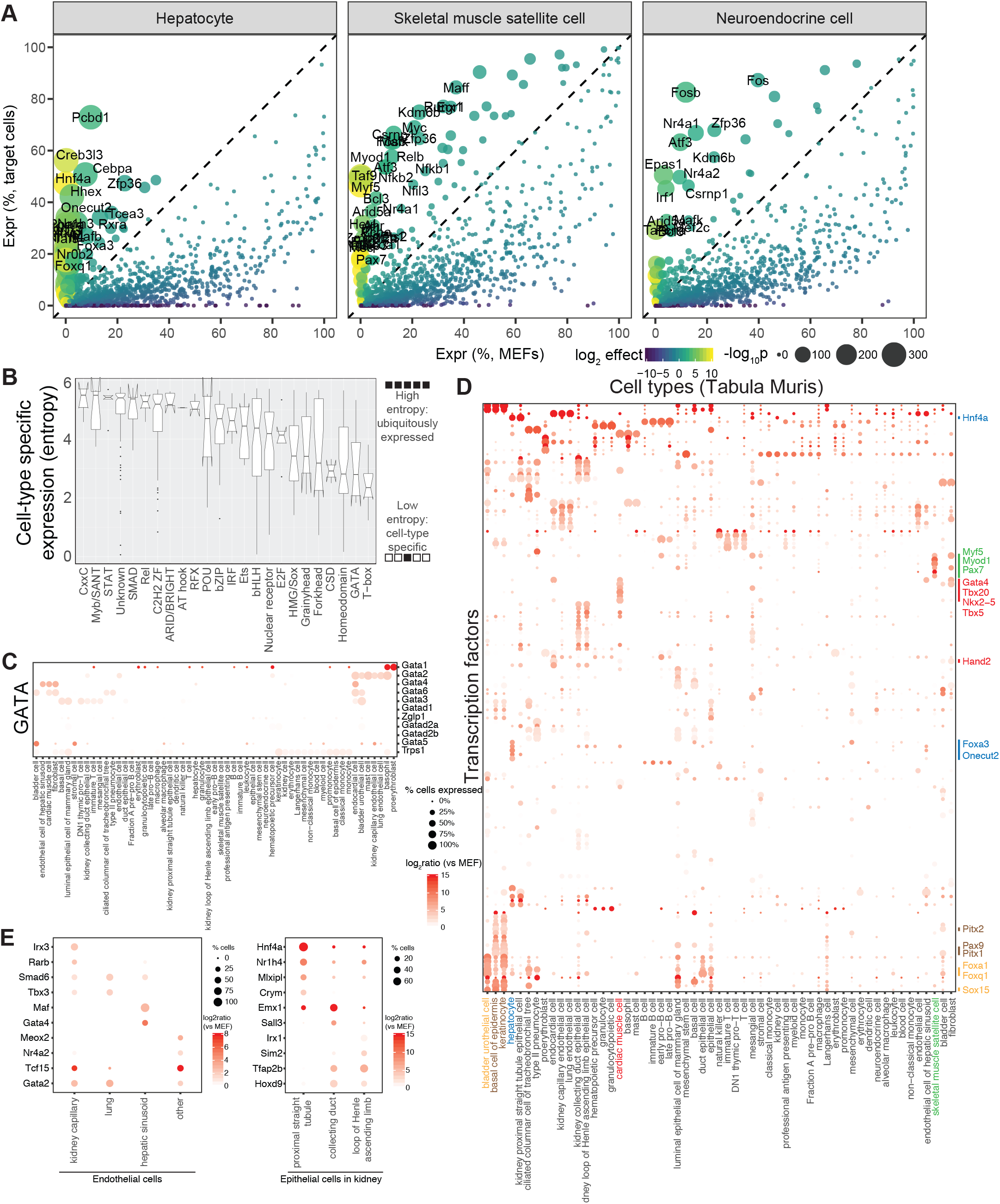
A roadmap for rational cellular reprogramming. A. Differential expression analysis of TFs in three primary mouse cell types (hepatocytes, skeletal muscle satellite cells, neuroendocrine cells) defined by Tabula Muris, as compared to MEFs. B. Boxplots indicating the cell-type specific expression of TF families based on Shannon entropy. C. Dot plot indicating the expression of 11 TFs from the GATA family across Tabula Muris cell types. D. Dot plot indicating the expression of 145 low entropy TFs across cell types defined by Tabula Muris. E. Dot plot of differentially expressed TFs for (left) endothelial cells across four mouse organs, and (right) three types of epithelial cells in mouse kidney.

To attain cell-type specificity, we reason that reprogramming cocktails will likely contain cell-type specifically expressed TFs. Using Shannon entropy (*32*), we observe that the GATA and T-box families are the most tissue-specific (**Figure 6B-C**), and many members have known roles in cellular reprogramming (*3*, *4*, *33*, *34*). Therefore, we used Shannon entropy to define 145 cell-type specifically expressed TFs across Tabula Muris cell types. Clustering of these cell-type specific TFs revealed several known TFs for direct reprogramming. For example, all of the known cardiomyocyte reprogramming factors (*Gata4*, *Hand2*, *Mef2c*, and *Tbx5*) are significantly enriched in cardiac muscle cells (**Figure 6D**, **S8**). Interestingly, Tabula Muris defines endothelial cells in four tissues, and we identify TFs specifically expressed in each one: kidney capillary (*Irx3*, *Rarb*), lung (*Smad6*, *Tbx3*), hepatic sinusoid (*Maf*, *Gata4*), and other tissue (*Meox2*, *Nr4a2*) (**Figure 6E**). In addition, Tabula Muris defines three types of epithelial cells in the kidney, each of which has cell-type specifically expressed TFs: proximal straight tubule (*Hnf4a*), collecting duct (*Emx1*), and loop of Henle ascending loop (*Irx1*). These unique TFs could have applications in cell-type specific, direct reprogramming.

## Discussion

Reprogram-Seq provides a functional platform for both undirected and directed cellular engineering. For undirected reprogramming, we provide two examples that highlight the robustness of Reprogram-Seq performed on 48F, which were curated based on their role in heart development and/or function from the literature. First, Reprogram-Seq accurately identified *MyoD1*-reprogrammed cells as induced skeletal myoblasts. Second, Reprogram-Seq specifically identified *Hand2* and *Gata6* as reprogramming factors for epicardial-like cells among a milieu of 48 factors. Alternatively, Reprogram-Seq can be used for rational reprogramming when a cell atlas is available. Here, Reprogram-Seq precisely identified 10 TFs that were subsequently refined to an optimized 3 TF combination capable of generating epicardial-like cells. Remarkably, *Hand2* and *Gata6* were identified by both undirected and directed reprogramming, thus confirming that both approaches achieve consistent solutions for generating epicardial-like cells and underscoring the robustness of Reprogram-Seq.

There are several advantages of Reprogram-Seq that distinguish it from previously described strategies for cellular engineering. First, Reprogram-Seq can be applied agnostic to literature. TF candidates for reprogramming can be derived entirely from cell-type specific single-cell atlases. In theory, each cell type expresses a unique profile of TFs, a subset of which are sufficient to establish the cell state. Second, as Reprogram-Seq can be applied in undirected reprogramming, it can be used for cellular engineering even if a single-cell atlas is not available for a particular cell type. Third, one Reprogram-Seq experiment can give insights into multiple TF cocktails. Thus, Reprogram-Seq is capable of providing refined TF cocktails, cell fate trajectories, and alternative cell fates by examining different subsets of cells from the same single-cell perturbation dataset.

Finally, Reprogram-Seq establishes an accelerated timeline of TF cocktail identification, optimization, and induced cell type characterization. To illustrate how Reprogram-Seq compares with traditional strategies for cellular engineering, we highlight induced cardiomyocyte (iCM) reprogramming as a well-studied example (*35*). The knowledge base for generating the original pool of candidate TFs for iCM reprogramming derived from many foundational studies using mouse genetics to understand heart development that spanned two decades (*36*). Then, the candidate pool was refined to GMT through an iterative screening process (*4*). Subsequent work identified additional factors to improve iCM reprogramming, identify alternative TF combinations, and further characterize the iCM phenotype (*35*). In addition, the trajectory of the early steps of iCM reprogramming was recently described using scRNA-seq (*37*). Taken together, these studies span more than twenty years of active investigation. Our example of applying Reprogram-seq for epicardial reprogramming significantly shortens this time frame. Thus, Reprogram-Seq is an important step towards acquiring the capability to reprogram new cell types for translational purposes on a reasonable time scale.

With the future completion of mammalian cell atlas projects (*27*, *38*), every cell type in human and mouse will be defined. This massive undertaking will provide a foundation for future mechanistic studies. We suggest that Reprogram-Seq can leverage this information to identify new TF cocktails for cell-type specific reprogramming. Reprogrammed human cells could then be made on demand and used for cell-based therapy, drug testing, and disease modeling to improve our understanding of human physiology.

## Author contributions

Conceptualization: N.V.M., and G.C.H.; Methodology: J.D., B.L., M.B., N.V.M., and G.C.H.; Software: J.D. and B.L.; Validation: M.B. and B.L.; Formal analysis: J.D. and B.L.; Investigation: J.D., M.B., B.L., P.Z., and S.X.; Writing -- Original Draft: J.D., B.L., N.V.M., and G.C.H.; Writing -- Review & Editing: J.D., B.L., N.V.M., and G.C.H.; Supervision: N.V.M. and G.C.H.; Funding acquisition: N.V.M. and G.C.H.

## Acknowledgements

We thank all the members in Hon and Munshi laboratories for insightful discussions and Ning Liu for reviewing the manuscript. This work is supported by the Cancer Prevention Research Institute of Texas (CPRIT) (RR140023 to G.C.H.), NIH (DP2GM128203 to G.C.H.; HL136604, HL133642, and HL133642 to N.V.M), the Department of Defense (PR172060 to G.C.H. and N.V.M.), the Welch Foundation (I-1926-20170325 to G.C.H.), the Burroughs Wellcome Fund (1009838 to N.V.M.), the March of Dimes Foundation (#5-FY13-203 to N.V.M.), and the Green Center for Reproductive Biology. We acknowledge the BioHPC computational infrastructure at UT Southwestern for providing HPC and storage resources that have contributed to the research results reported within this paper. We also acknowledge UT Southwestern’s McDermott Center for providing next-generation sequencing services for this work.

## SUPPLEMENTAL TEXT

### Additional analysis of Cluster 20 cells from Figure 1

On average, cells in Cluster 20 express 61.3 times more *MyoD1* as compared to other reprogrammed MEFs (**Figure 1D-E**) (Cluster 20: 254 cpm; other reprogrammed MEFs: 4.1 cpm; p < 2.2e-16; Wilcoxon test). In addition, 164 of the 165 cells in Cluster 20 are reprogrammed cells, and none belong to uninfected MEFs. Cluster 20 exhibits both exogenous as well as endogenous expression of *MyoD1* (**Figure S3A**).

### Additional analysis supporting 3F as an optimized reprogramming cocktail

First, we examined the expression of the transcription factors in the epicardial-like Cluster 10. While each of the factors in 10F exhibited some induction, we observed dramatic enrichment of *Atf3*, *Gata6*, and *Hand2* expression in the reprogrammed cells of Cluster 10, with over half of the cells expressing these factors (**Figure S5A-B**). We denote this combination of TFs as 3F. Collectively, 3F is >8.1-fold more enriched in Cluster 10 reprogrammed cells compared to uninfected MEFs, and > 2.6-fold more enriched compared with other reprogrammed MEFs. Supporting the consistency of Reprogram-Seq, *Gata6* and *Hand2* were also identified in our unbiased analysis of 48F above.

Next, we sought to confirm that reprogrammed cells in Cluster 10 have exogenous expression of 3F, as we reasoned that enrichment of exogenously expressed trans-genes is indicative of successful viral induction. We took advantage of our cloning strategy in that exogenous TFs do not have 3’UTRs. Therefore, reads derived from exogenous TFs will be strictly upstream of the 3’UTR while reads derived from endogenous TFs will be enriched downstream of the 3’UTR. In primary epicardial cells, we observed that 92.0% of reads for *Hand2* fall into the 3’UTR, which is consistent with endogenous expression. In contrast, this number is only 2.5% for cells in Cluster 10, which is indicative of exogenous expression (**Figure S5C**). Control genes including *Gapdh* and *Bnc1* are almost exclusively endogenously expressed (**Figure S5C**). By using this observation to estimate the exogenous and endogenous expression of 10F, we confirmed that cells in Cluster 10 exhibit significant exogenous expression of 3F (**Figure 3A**). We also examined whether cells with 3F expression are more likely to be in Cluster 10. Of the 1026 MEF-derived cells expressing 3F, we observe that 555 (54.1%) belong to Cluster 10 as compared to 18.4% expected by chance (p = 4.10e-197). These results suggest that 3F may be an optimized TF cocktail for epicardial reprogramming.

### Additional analysis of alternative cell fates during 3F reprogramming

Pseudotime also identifies two states (E and G) that branch off the main trajectory. These cells exhibit expression of 3F (**Figure S6A,C**), indicating that they are indeed derived from MEF reprogramming. However, the expression of endogenous epicardial markers is lower in these cells compared to the terminal state H (**Figure S6C**). These observations suggest that states E and G could be alternative cell fates of 3F reprogramming. Next, we examine potential molecular pathways interfering with epicardial reprogramming. We observe that cells in state E exhibited higher expression of cell division genes (**Figure S6D**) (p = 2.4E-11) including the DNA topoisomerase *Top2a*, the cytokinesis regulator *Prc1*, and the mitotic regulator *Cdca8* (**Figure S6E**). Consistent with previous studies (*1*–*4*), these results suggest that inhibiting the cell cycle could improve epicardial reprogramming. Finally, extending this analysis to state G, we find that this alternative cell fate exhibits failed activation of several tyrosine kinases with roles in cell signalling, including *Abl2*, *Ptk2*, and *Src*. Consistent with this observation, gene ontology analysis reveals that genes repressed in state G are enriched for small GTPase mediated signal transduction (p = 1.2E-09) and epidermal growth factor receptor signaling pathway (p = 9.5e-06) (**Figure S6B**). These results suggest that activation of specific signal transduction pathways may be an important step in epicardial reprogramming.

## KEY RESOURCES TABLE

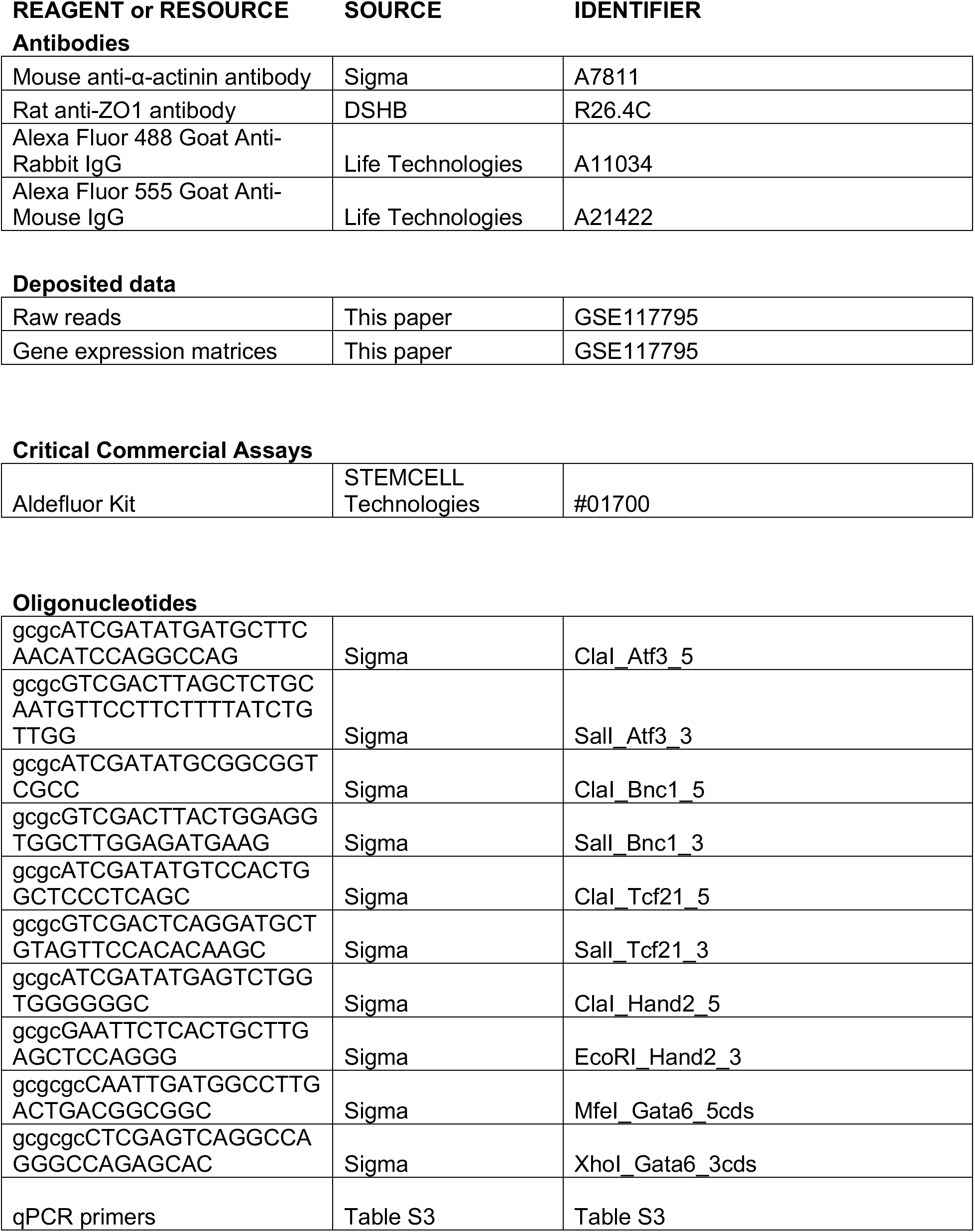

Software and Algorithms
|  |  |  |
| --- | --- | --- |
| bcl2fastq (version 2.17.1.14) | Illumina | https://support.illumina.com/sequencing/sequencing_software/bcl2fastq-conversion-software.html |
| Cell Ranger (version 2.1.0) | 10x Genomics | https://www.10xgenomics.com |
| Drop-seq Tools (version 1.12) | Macosko et al (2015) | http://www.dropseq.org |
| featureCounts (version 1.5.3) | Liao et al., (2014) | http://bioinf.wehi.edu.au/featureCounts |
| Gviz (version 1.24.0) | Hahne and Ivanek (2016) | https://bioconductor.org/packages/release/bioc/html/Gviz.html |
| irlba (version 2.3.2) | Jim Baglama, Lothar Reichel, B. W. Lewis | https://cran.r-project.org/web/packages/irlba/index.html |
| Monocle (version 2.8.0) | Qiu et al., (2017) | http://cole-trapnell-lab.github.io/monocle-release/ |
| R (version 3.5.1) | R Foundation | https://www.r-project.org |
| Rtsne (version 0.13) | van der Maaten (2014) | https://cran.r-project.org/web/packages/Rtsne/index.html |
| SAMtools (version 1.8) |  | https://www.htslib.org |
| STAR (version 2.5.3a) | Dobin et al., (2013) | https://github.com/alexdobin/STAR |
| sva (version 3.28.0) | Johnson et al., (2007) | https://bioconductor.org/packages/release/bioc/html/sva.html |
| topGO (version 2.32.0) | Adrian Alexa, Jorg Rahnenfuhrer | https://bioconductor.org/packages/release/bioc/html/topGO.html |
| UMI-tools (version 0.5.3) | Smith et al., (2017) | https://github.com/CGATOxford/UMI-tools |
| org.Mm.eg.db (version 3.4.1) | Marc Carlson | https://bioconductor.org/packages/release/data/annotation/html/org.Mm.eg.db.html |
| GO.db (version 3.4.1) | Marc Carlson | https://bioconductor.org/packages/release/data/annotation/html/GO.db.html |
| igraph (version 1.2.1) | https://cran.r-project.org/web/packages/igraph/AUTHORS | http://igraph.org/ |
| RANN (version 2.6) | Sunil Arya and David Mount (for ANN), Samuel E. Kemp, Gregory Jefferis | https://cran.r-project.org/web/packages/RANN/index.html |
| Code for this manuscript | This paper | https://github.com/jlduan/Reprogram-Seq |

## METHOD DETAILS

### Primary Sample Collection

Pregnant dams were housed in a standard facility 12hr light/dark cycle, 70-75°F, and ∼50% humidity. Primary cardiac cells were harvested from neonatal mouse hearts at P0 by dissecting and mincing the hearts. The tissue was then incubated in trypsin for a total of 30 minutes in a shaking water bath at 37°C. Supernatant was strained and transferred to a flask containing cold growth media every 5 minutes-6 fractions total. Cells were spun at 500X G for 5 minutes at 4°C and resuspended in cold growth media. Incubation times in trypsin were adjusted depending on the dissected area of interest.

### Construction of Retroviral Vectors

Retroviral vectors were generated by subcloning individual TF coding sequences into the pBABE plasmid (1). TF coding sequences were amplified from mouse or human heart cDNA using primers containing linkers with appropriate restriction enzyme sites. Amplified PCR products were cut with restriction enzymes, gel purified, and ligated into the pBABE vector. Positive clones were verified by Sanger sequencing.

### MEF Isolation

Mouse embryonic fibroblasts (MEFs) were isolated from timed pregnant E12 CD-1 mice purchased from Charles River as described previously (1). Briefly, uterine horns containing embryos were removed from the pregnant dam and placed in cold DPBS. Subsequently, embryos were removed from the uterus and amniotic sac. Head, limbs, tail, and internal organs and viscera were removed and discarded. Remaining tissue was minced, washed with DPBS, and incubated in trypsin 0.25% for 10 minutes at 37°C. Cells were strained using 100µm filter. Trypsin was inactivated with growth media (high glucose DMEM, 10% FBS, 1X penicillin/streptomycin). Cells were spun at 500X G for 5 minutes at 4°C and resuspended in growth media and plated at ∼100K cells per cm^2^. MEF’s were maintained in growth media and incubated at 37°C in 5% CO_2_. After 24 hours, cells were frozen at -80°C in aliquots of 5-10 million cells suspended in 90% FBS and 10% DMSO.

### Viral Packaging, Infection, and Single Cell Reprogramming

A packaging cell line Platinum-E (Cell Biolabs, San Diego, CA) was used to produce retrovirus for reprogramming (1). Packaging cells were transfected using Fugene (Promega, Madison, WI) per manufacturers’ instructions with retroviral vectors (pBabe) expressing specific transcription factors. Virus was harvested, filtered (0.45µm pore) and applied to MEFs at 24 hrs and 48 hrs post-transfection after the addition of polybrene 8µg/mL (Sigma-Aldrich, St. Louis, MO). The cells were incubated for 7 or 14 days and subsequently washed with DPBS and trypsinized into a single cell suspension. Cells were spun at 500X G for 5 minutes at 4°C and resuspended in cold growth media.

### Phase Contrast Microscopy

For phase contrast microscopy, cells were imaged live in media on a 24-well plate with EVOS™ FL Auto Imaging System (Thermo Fisher Scientific, Waltham, MA). Phase contrast mode was used for imaging.

### Cell Staining

For ICC staining, MEFs were plated on 12mm glass coverslips #1.5 for 16-18 hrs then infected with retrovirus. After 7 days, cells were washed with PBS and fixed using 4% Paraformaldehyde (Electron Microscopy Sciences, Hatfield, PA) for 15 minutes at room temperature. Cells were washed 3 times with PBS then permeabilized using PBST (0.1% TritonX-100 in PBS) 3 times for 5 minutes. Cells were then blocked for 10 minutes at room temperature using Universal Blocking Buffer diluted to 1x. Primary antibodies were diluted in 1x Universal Blocking Buffer— ZO1(1:5)(DSHB), a-actinin (1:200) (Sigma-Aldrich, St. Louis, MO)—and incubated overnight at 4°C after application to the cells. Cells were then washed 3 times with PBST and incubated with secondary antibody for 1 hr at room temperature. All secondary antibodies were obtained from Life Technologies’ (Carlsbad, CA) Alexa Fluor Dyes 488, 555, and 647. Cells were washed 3 times and coverslips were mounted using Vectashield with DAPI (Vector Laboratories, Burlingame, CA) and imaged using a Nikon A1R confocal microscope.

### Aldefluor Assay

MEFs were incubated for 7 days and 14 days post induction to assess aldehyde dehydrogenase activity (STEMCELL Technologies, Vancouver, Canada). Aldefluor staining was carried out according to the manufacturers’ directions using an incubation time of 50 minutes at 37°C. Flow cytometry was carried out on a BD FACSCalibur flow cytometer (Becton Dickinson, Franklin Lakes, NJ) and control signals was subtracted from each sample.

### Single Cell RNA Sequencing Library Preparation

To confirm the single cell resolution of our RNA sequencing, prior to real experiments, mixed-species experiments were performed. Briefly, human cell lines and mouse primary/reprogrammed cells were mixed together and scRNA-seq libraries were constructed using the Drop-seq and 10x Genomics platforms. The resulting data of both platforms are highly organism-specific, suggesting a low cell multiplet rate and low cross-cell contamination.

#### Drop-seq Procedure

Drop-seq was performed as described previously (*2*, *3*) with small modifications. Briefly, droplets were generated using microfluidic devices, which encapsulated single cells and barcoded beads (ChemGenes Corporation, Wilmington, MA, catalog number Macosko201110). Beads were collected after the breakage of individual droplets using perfluorooctanol (Sigma-Aldrich, St. Louis, MO, catalog number 370533). RNA hybridized on the barcoded beads was reverse transcribed using Maxima H minus reverse transcriptase (Thermo Fisher Scientific, Waltham, MA, catalog number EP0751) and cDNA was amplified with 13 cycles (KAPA HotStart ReadyMix, Kapa Biosystems, Wilmington, MA, catalog number KK2602). In-house Tn5 transposase was used to insert sequencing adapters and fragmentate cDNA. Illumina Nextera adapters i5 and i7 were used to amplify the fragmented cDNA with 12 cycles (KAPA HiFi PCR Kits, catalog number KK2102). Agarose gel size selection was performed to recover fragments with length of 400 to 600 bp.

#### 10x Genomics Single Cell RNA Sequencing Procedure

The concentration of single cell suspension was adjusted to 500-1000 cells/µL and was loaded on the 10x Genomics Chromium™ system (10x Genomics, Pleasanton, CA) with the aim of generating 6000 to 10000 transcriptomes per channel (Chromium™ Single Cell 3’ Library & Gel Bead Kit v2, catalog number 120237). Single cell RNA sequencing libraries were constructed following the manufacturer’s instructions (4).

## QUANTIFICATION AND STATISTICAL ANALYSIS

### Sequencing, Basecalling and Demultiplexing

Libraries were sequenced on Illumina NextSeq 500/550 sequencing systems (Illumina, San Diego, CA). BCL files generated by Illumina sequencing systems were demultiplexed and converted to standard FASTQ files using bcl2fastq (version 2.17.1.14). For 10x Genomics libraries, the mkfastq function from Cell Ranger pipeline (version 2.1.0) was used to perform basecalling and demultiplexing for downstream analyses.

### Read Alignment and Generation of Gene Expression Matrix

For Drop-seq data, demultiplexed reads were processed using Drop-seq Tools (version 1.12) as described in Macosko et al. (2) with small modifications. Briefly, low quality reads containing cell barcodes and UMIs were filtered and polyA tails (longer than 6 bases) were trimmed before mapping. Processed reads were mapped to mouse mm10 reference using STAR (version 2.5.3a) (5). Only uniquely aligned reads were kept. Cell barcodes within one edit distance were collapsed together. UMI-based PCR duplicate removal was performed using the directional method of UMI-tools with default parameters (version 0.5.3) (6). An inflection point of cumulative number of reads of cells in each library was calculated to determine the number of cells in each experiment. Gene expression matrices were generated using featureCounts (version 1.5.3) (7). The gene annotation used is based on Ensembl release 84 (GENCODE Gene Set M9).

For 10x Genomics data, the gene expression matrix was generated for each experiment using Cell Ranger pipeline (version 2.1.0) with default parameters, except the parameter of expected number of cells, which was adjusted based on each individual experiment. The genome reference and gene annotation used were the same as Drop-seq data.

### Dimensionality Reduction

#### Filtering Low Quality Cells and Uninformative Genes

All Drop-seq and 10x Genomics data were combined together respectively and analyzed separately. Cells with fewer than 200 genes detected were filtered. Genes detected in fewer than 30 cells or with fewer than 60 UMIs in total across all cells were filtered. In total, 25,776 cells from Drop-seq and 34,564 cells from 10x Genomics data were kept for downstream analyses.

#### Normalization and Batch Correction

The total UMI counts for each cell were normalized towards the median UMI counts per cell by multiplying each cell with a scaling factor. The resulting matrix was natural-log-transformed with the addition of a pseudocount of 1 prior taking the log-transformation. The ComBat (8) method from sva package (version 3.28.0) was used to correct the batch effect with the default parametric adjustment parameters (9).

#### Principal Component Analysis (PCA)

The median-normalized natural-log-transformed batch-effect-corrected matrix was centered and scaled per gene so that the mean expression value of each gene is 0 and the standard deviation is 1 across all cells. The resulting matrix was used to perform PCA using the prcomp function from built-in R package stats (version 3.5.1). To determine the number of principal components to use for clustering and visualization, permutation tests were performed (2). PCA was performed on 500 permuted versions of the original matrix. In each version, 10% of the total genes were randomly selected and their expression values were independently permuted. To speed up the calculation, prcomp_irlba function from irlba package (version 2.3.2) was used to perform PCA on the original and 500 permuted matrices, and only the top 40 principal component vectors were calculated. The average proportions of variance explained by each principal component of the 500 permutations were compared to the proportions of variance explained by each principal component of the original matrix.

#### t-Distributed Stochastic Neighbor Embedding (t-SNE) Visualization

A two-dimensional non-linear embedding of the single cells was computed using Barnes-Hut implementation of t-SNE method (10). The scores of the significant principal components estimated above (output of the prcomp function) were used as input to Rtsne package (version 0.13). The initial PCA step was disabled, and perplexity parameter was set to 30 with 3000 iterations.

### Graph Clustering

The Louvain-Jaccard method (11) was used to cluster single cells. The input is the scores of the chosen principal components. Briefly, a *k*-nearest neighbor (k-NN) graph was built based on euclidean distance in chosen principal component scores using nn2 function from RANN package (version 2.6, *k* set to 30). Jaccard overlap index of each edge in the graph was calculated and used as input to the Louvain algorithm implemented in igraph package (version 1.2.1).

### Binomial Test for Differentially Expressed Genes

The method used for testing if a gene is expressed more frequently between different group of cells was described in Shekhar et al. (11)

### Identification of Exogenous and Endogenous Expression

The plasmid sequences of Myrf Sox6 Tbx20 and Wt1 are from human. To accurately calculate their expression values, demultiplexed reads were also mapped to human GRCh38 reference. Reads uniquely mapped to mouse genomic regions of these four genes and reads uniquely mapped to the orthologs of those fours gene in human genome were retrieved. The source of each read was determined based on their genotypes.

Genomic tracks were generated using uniquely mapped non-PCR duplicate reads from selected group of cells. The distribution pattern of reads were compared among different cells groups. The gene sequences on the retroviral plasmids have no 3’ untranslated regions (3’ UTRs).

### Pseudotime Construction

Monocle (version 2.8.0) was used for pseudotime analysis of single cells (12). Raw UMI counts were used as input and the distribution was set to “negbinomial.size()”. The minimum expression level that constitutes true expression was set to 0.5. Differentially expressed genes between starting cells and target primary cells were used to measure cells’ progresses during reprogramming. The data dimensionality was reduce to 2 using “DDRTree” method.

### Gene Ontology (GO) Enrichment Analysis

Enrichment analysis for GO terms was performed using topGO (version 2.32.0). Fisher’s exact test was used for enrichment tests.

### Transcription Factor (TF) Annotation

TFs were defined based on GO terms (org.Mm.eg.db, version 3.4.1; GO.db, version 3.4.1). Genes need to meet both criterias to be considered as TFs: a) have either “regulation of transcription” or “transcription factor activity” GO terms in BP sub-ontology; b) “DNA binding” or “transcription factor activity” GO terms in MF sub-ontology.

### Transcription Factor Family Categorization

A full list of curated human TFs was obtained from the “HumanTFs” website (http://humantfs.ccbr.utoronto.ca/) (13). The identification of TF families was based on the third column (DNA-binding domain, or DBD) of the list. Next, the mouse orthologs of all curated human TFs were identified through an online tool, bioDBnet (https://biodbnetabcc.ncifcrf.gov/db/dbOrtho.php). Human Ensembl IDs were converted to mouse gene symbols, and the generated list was overlapped with the list of all annotated mouse TFs mentioned above. Only the common TFs between the two lists were used for downstream analysis.

## DATA AND SOFTWARE AVAILABILITY

The single cell RNA sequencing data reported in this paper were deposited in NCBI Gene Expression Omnibus (GEO) under the accession number GSE117795. Scripts for analyzing Reprogram-Seq data is available at https://github.com/jlduan/Reprogram-Seq.

## SUPPLEMENTAL FIGURE LEGENDS

**Supplemental Figure 1:**
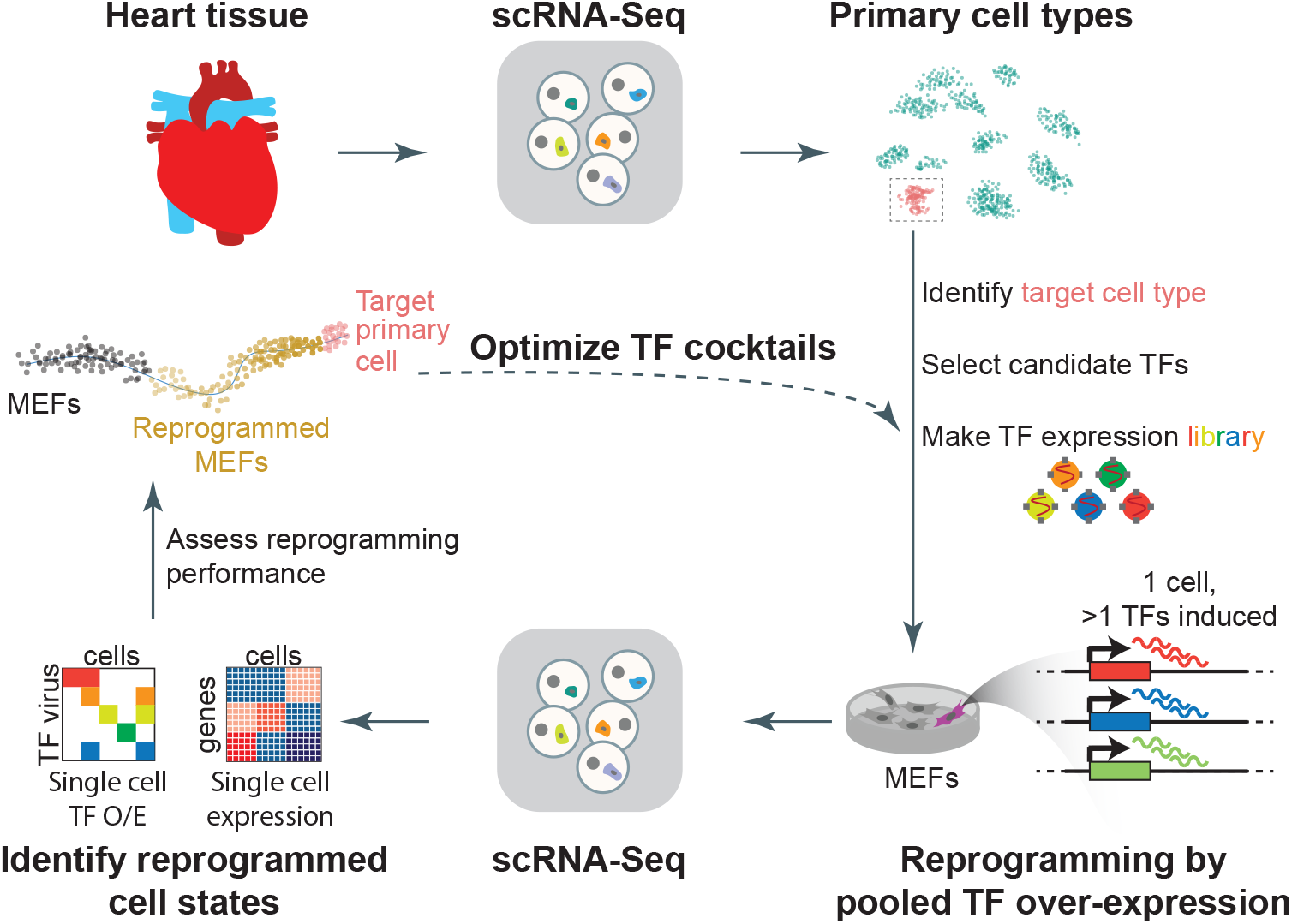
Combinatorial cellular reprogramming with Reprogram-Seq. Analysis of single-cell transcriptomes or expert knowledge identifies a set of candidate TFs for direct reprogramming. TFs are cloned into retroviral expression vectors and induced in MEFs. In Reprogram-Seq, we perform scRNA-Seq on thousands of individual cells to identify successfully reprogrammed cells and the exogenous factors expressed in these cells. The process can then be repeated with these factors to optimize TF cocktails for direct cellular reprogramming.

**Supplemental Figure 2:**
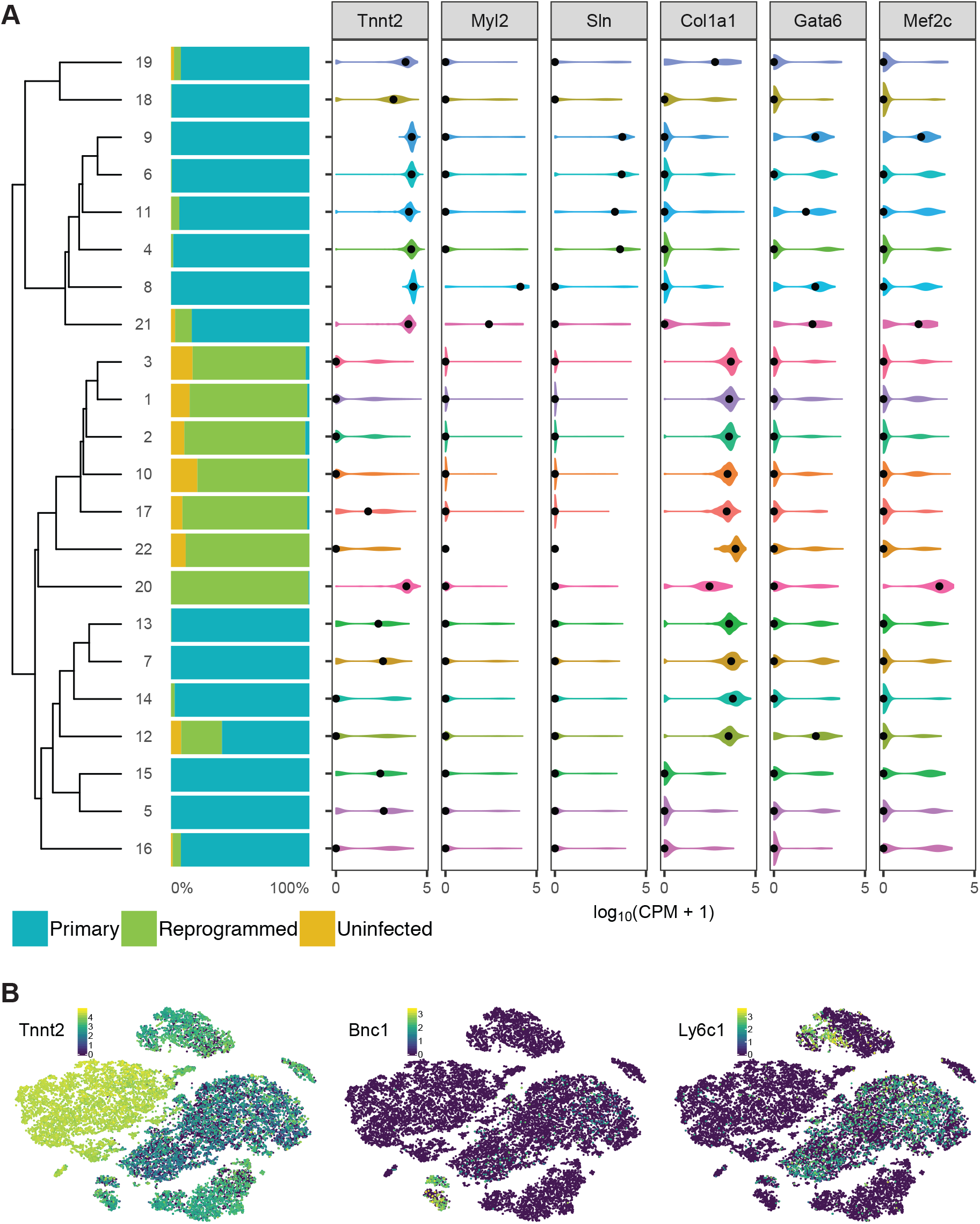
A. (left) Bar charts indicating the cellar composition of t-SNE clusters defined in Figure 1B. (right) Violin plots illustrating the expression of cardiac markers from single-cell expression data derived from P0 mouse heart and reprogrammed/uninfected MEFs. B. Heatmap of cells in Figure 1B by the expression of (left) the cardiomyocyte marker gene *Tnnt2*, (middle) the epicardial marker gene *Bnc1*, and (right) the lymphocytic marker gene *Ly6c1*.

**Supplemental Figure 3:**
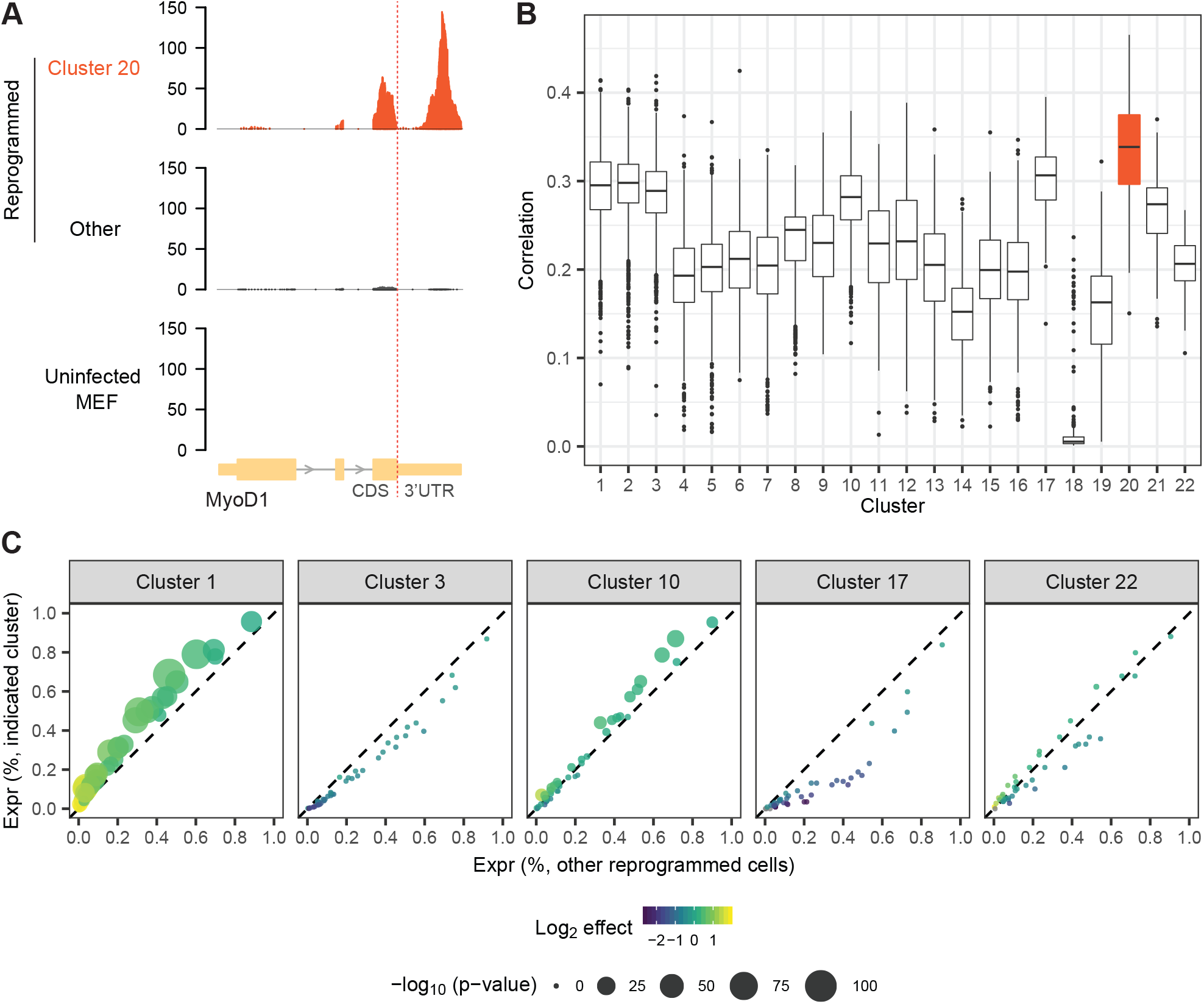
A. Genome Browser snapshot of MyoD1 expression from Cluster 20 cells, other reprogrammed cells, or uninfected MEFs. The cloned exogenous MyoD1 gene does not have a 3’ UTR. Hence, exogenous reads are concentrated upstream of the 3’ UTR, while endogenous reads are inside the 3’ UTR. B. We performed scRNA-Seq on MyoD1-reprogrammed MEFs. Shown is the gene expression correlation of MyoD1-reprogrammed MEFs with the clusters from Figure 1B. C. Differential expression analysis of 48F in MEF-derived clusters, as compared to all other reprogrammed cells. Each plot contains 48 dots (colored by fold change and sized by p-value).

**Supplemental Figure 4:**
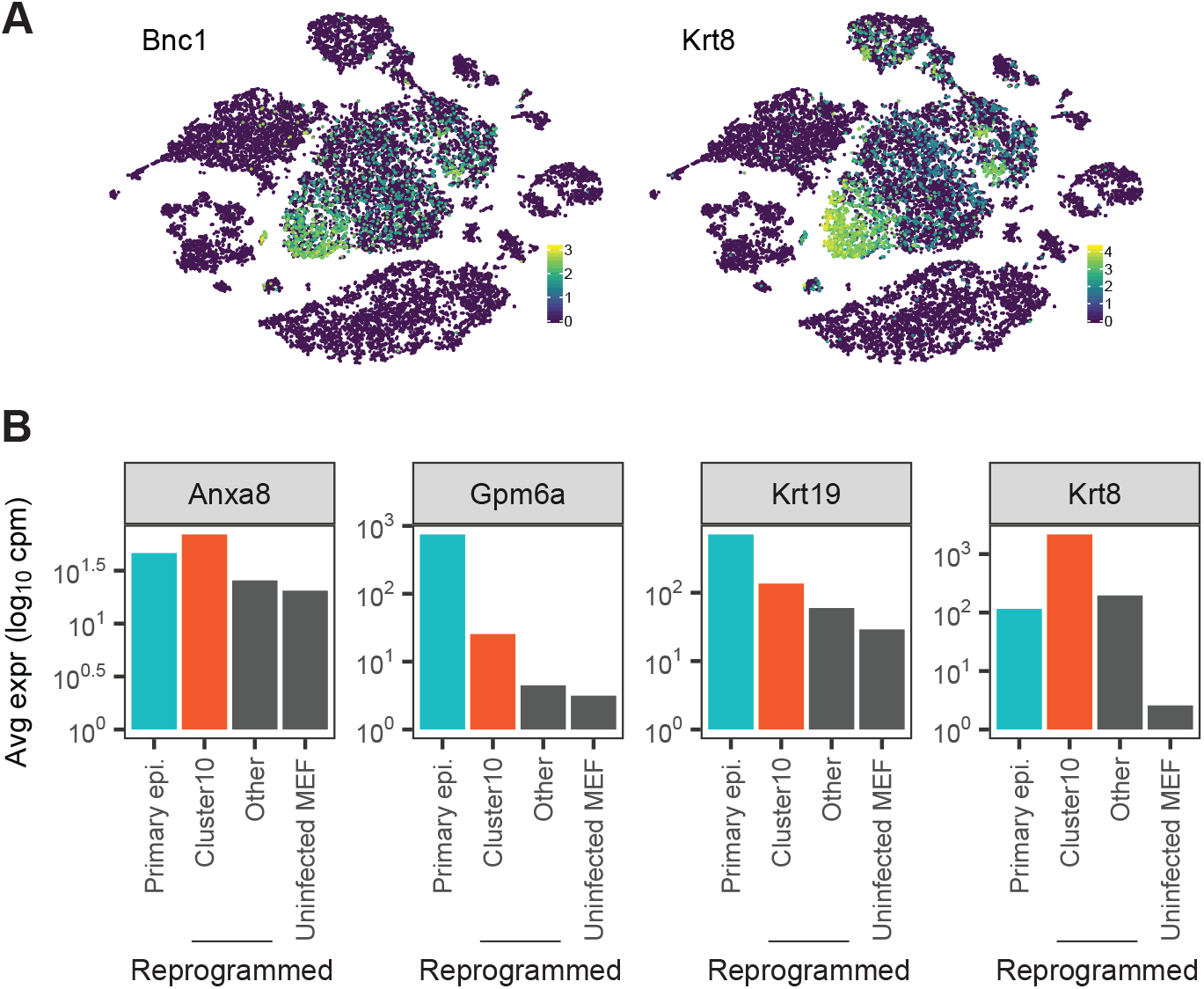
A. Heatmap of cells in Figure 2C by the expression of (left) the epicardial marker genes *Bnc1* and *Krt8*. B. Expression of epicardial marker genes *Anxa8*, *Gpm6a*, *Krt19*, and *Krt8* in primary and MEF-derived cells.

**Supplemental Figure 5:**
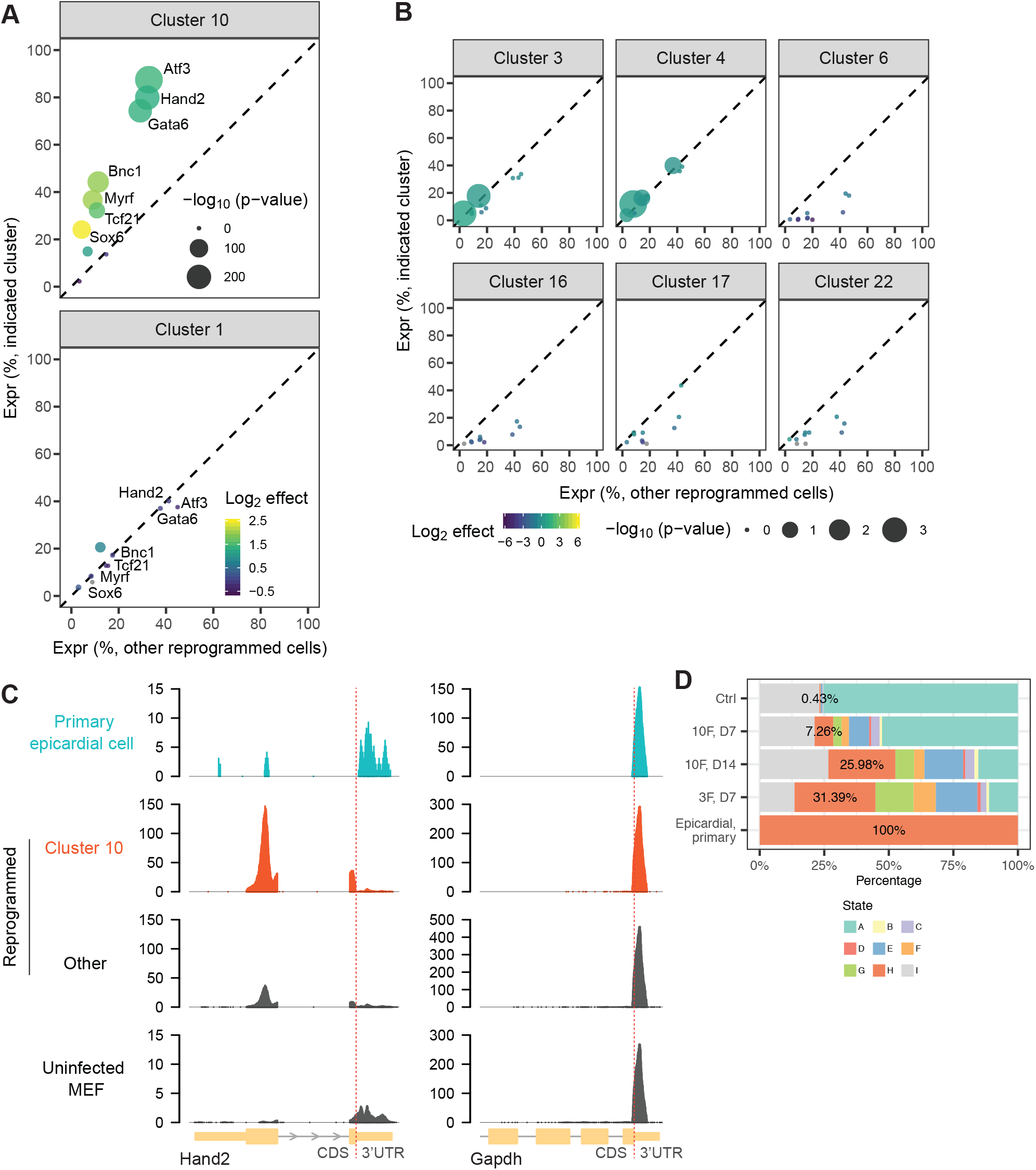
A-B. Differential expression analysis of 10F in MEF-derived clusters, as compared to all other reprogrammed cells. C. Genome browser snapshots of (left) Hand2 and (right) Gapdh. In primary cells and uninfected cells, Hand2 reads are almost exclusively derived from the 3’UTR. In contrast, read enrichment in reprogrammed cells is rarely in the 3’ UTR, which is indicative of exogenous expression. D. Bar plot indicating the composition of cells in each reprogramming experiment based on pseudotime states from Figure 3C.

**Supplemental Figure 6:**
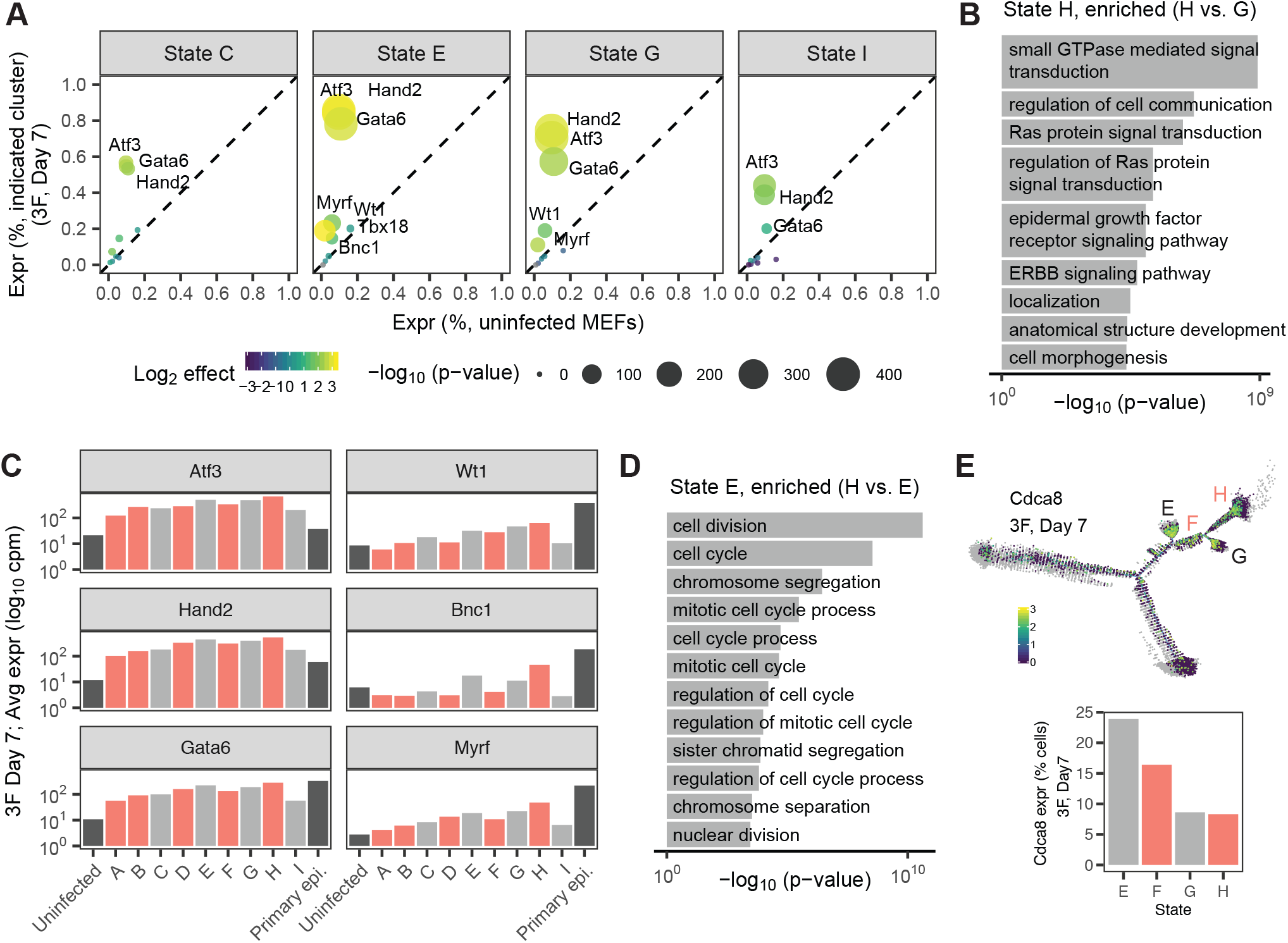
A. Expression of exogenous genes (*Atf3*, *Hand2*, *Gata6*) and endogenous genes (*Myrf*, *Wt1*, *Bnc1*) across pseudotime states in MEFs reprogrammed with 3F for 7 days. B. Gene ontology analysis of genes differentially expressed in psuedotime state G. C. Bar plot indicating the expression of exogenous genes (*Atf3*, *Hand2*, *Gata6*) and endogenous genes (*Myrf*, *Wt1*, *Bnc1*) across pseudotime. D. Gene ontology analysis of genes differentially expressed in psuedotime state E. E. (top) Heatmap of cells by the expression of the cell division marker *Cdca8*. (bottom) Quantification of *Cdca8* expression in pseudotime states E-H.

**Supplemental Figure 7:**
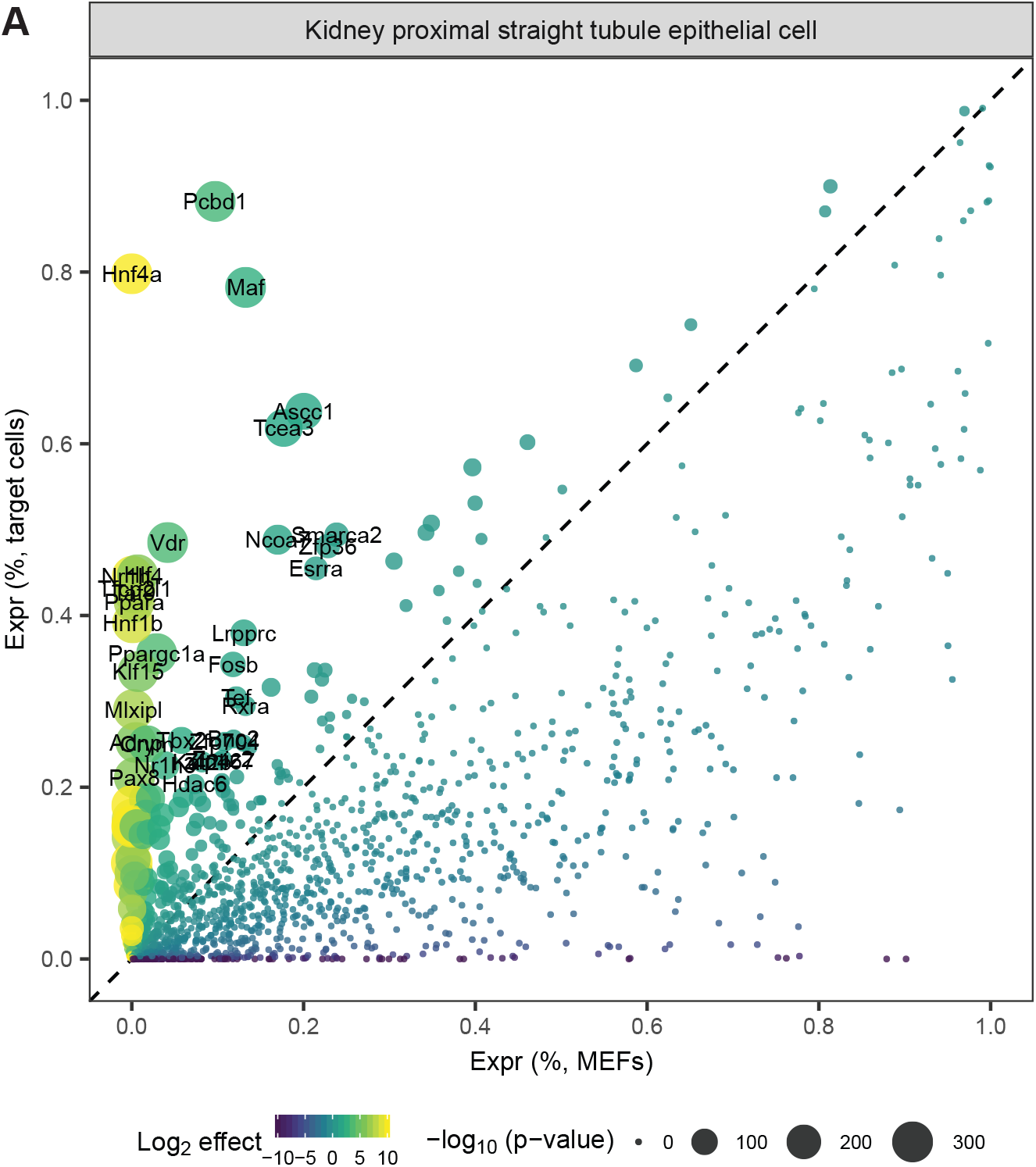
A. Differential expression analysis of TFs in renal tubular epithelial cells defined by Tabula Muris, as compared to MEFs.

**Supplemental Figure 8:**
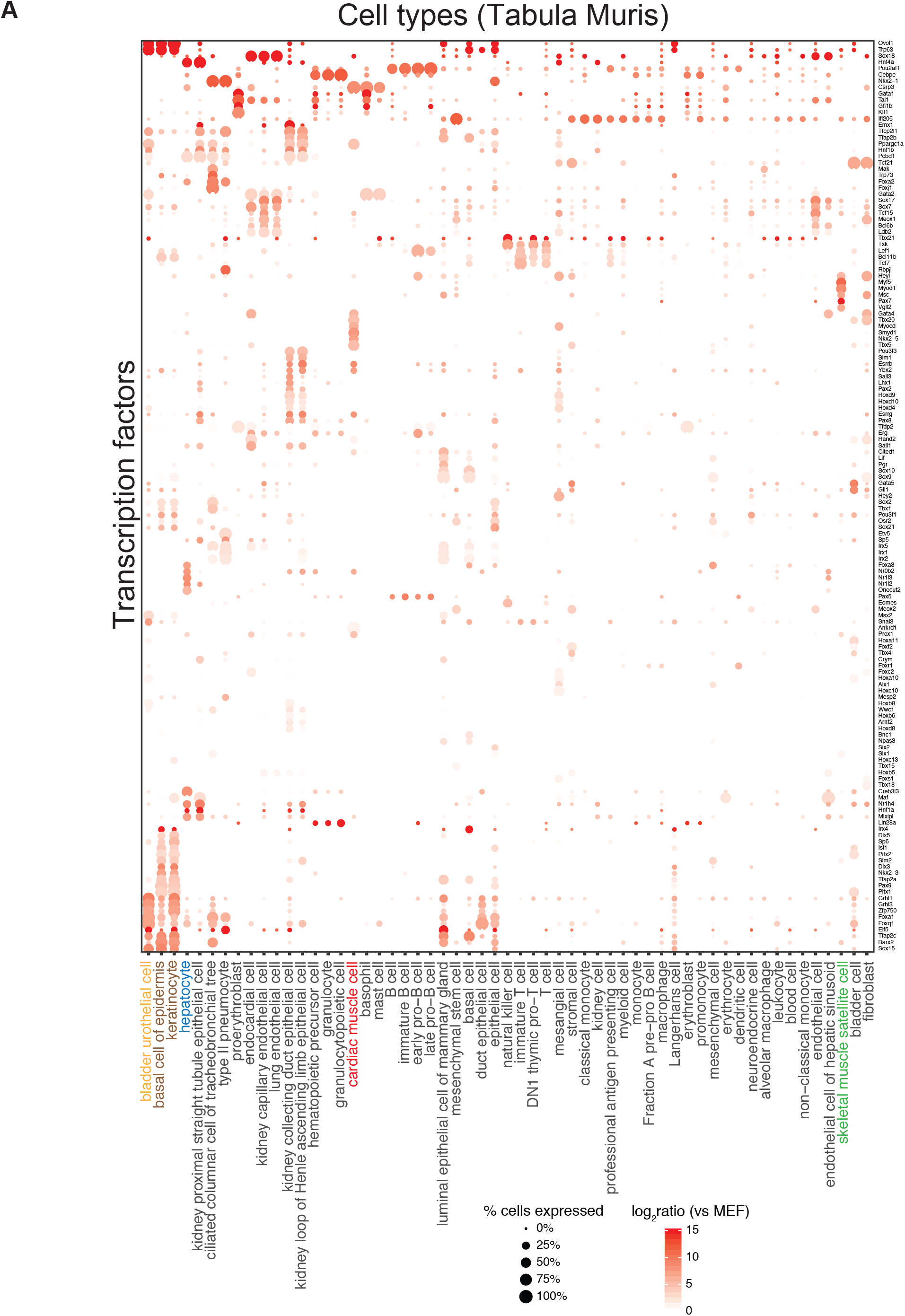
A. Dot plot indicating the expression of 145 low entropy TFs across cell types defined by Tabula Muris. This is an enlarged version of Figure 6D.

## SUPPLEMENTAL TABLE LEGENDS

**Table S1: Sequencing statistics.**

**Table S2: Transcription factors and genes in 48F.**

**Table S3: qPCR primers.**

**Table S4: Top differentially expressed TFs for every cell type identified by Tabula Muris Consortium.**

